# The meiotic synaptonemal complex is assembled from independently regulated gene programmes

**DOI:** 10.64898/2026.09.09.750351

**Authors:** LC Brannan, CJ Fawcett, IF Sou, AD Bates, A Saifudeen, MA Ikert, AG Ginesta, B Hammond, UL McClurg

## Abstract

Synaptonemal complex (SC) proteins assemble a highly specialised chromosome structure during meiosis and are considered products of a coordinated germline programme that is silenced in somatic cells. Here, we show that this binary model does not describe the regulation of the eight core mammalian SC genes. Despite assembling into a single molecular complex, SC genes follow distinct regulatory trajectories from meiotic entry through SC disassembly. Their activation is staggered, transcripts and proteins persist with different kinetics after SC disassembly begins in late pachytene, and RNA abundance generally fails to predict protein abundance, revealing extensive regulation between transcription and protein accumulation. Integrating promoter state, nascent transcription, RNA and translational measurements identified gene-specific regulatory strategies rather than a shared SC regulatory mechanism. This independence extends beyond the germline and explains how cancers can express individual SC genes. In cancer cells, individual SC loci occupy distinct transcriptional states, including conventional promoter activation, alternative promoter usage, cell-cycle-dependent transcription and promoter competence without detectable productive transcription. DNA methylation can repress individual SC promoters but does not define a common somatic OFF state, while related transcriptional inputs produce different downstream RNA outputs between SC genes. Unexpectedly, single-cell transcriptomes across healthy mouse and human tissues reveal that somatic SC expression is not restricted to cancer: individual SC genes show reproducible associations with specific cell populations, including fibroblasts, myeloid cells, Schwann cells and progenitors. Thus, SC proteins are neither expressed nor silenced as an obligately coupled gene set. We propose that SC identity emerges from the temporally restricted convergence and assembly of independently regulated genes and proteins during meiosis, while their regulatory autonomy permits individual components to be retained or redeployed in normal somatic cells and cancer. The SC therefore represents an emergent molecular state: its components retain distinct regulatory identities, while their transient convergence during meiotic prophase generates a structure and function that none defines individually.

## Introduction

The assembly of large macromolecular structures requires cells to produce multiple protein components in the correct amounts, cellular locations and developmental windows. This creates a fundamental regulatory problem: proteins that function together may be encoded by genes located at independent genomic loci and subject to distinct transcriptional and post-transcriptional controls. Coordinated expression is therefore often assumed to be an important feature of molecular-complex biogenesis. Conversely, when a specialised structure is no longer required, its components might be expected to undergo coordinated transcriptional silencing. However, whether the genes encoding highly stage-restricted cellular structures are controlled in this binary, modular manner remains poorly understood.

The synaptonemal complex (SC) provides a powerful system to address this question. The SC is a large, meiosis-specific protein structure assembled between homologous chromosomes during meiotic prophase I (1). Its core architecture in mammals is constructed from a small and well-defined set of proteins, including the chromosome-axis components SYCP2 (2) and SYCP3 (3), the transverse-filament protein SYCP1 (4), and the central-element proteins SYCE1, SYCE2 (5), SYCE3 (6), TEX12 (7) and SIX6OS1 (8). Together, these proteins generate a highly ordered structure that promotes homolog alignment and supports the progression and regulation of meiotic recombination. Following pachytene, the SC is disassembled as homologous chromosomes desynapse (9), providing an unusually clear biological transition between a developmental state in which the complete molecular structure is required and one in which it is not.

SC proteins have consequently been regarded as products of a tightly restricted meiotic gene-expression programme (1). This view is consistent with the profound meiotic defects caused by disruption of individual SC components (3, 5, 6, 8, 10) and with their historically described enrichment in germ cells. More broadly, germline-restricted genes are frequently conceptualised as existing in binary transcriptional states: activated during germ-cell differentiation and epigenetically repressed in somatic tissues. DNA methylation has featured prominently in models explaining this restriction, particularly because aberrant promoter hypomethylation can accompany expression of germline and cancer-testis genes in tumours (11–13). Such models predict that genes encoding components of the same meiosis-specific structure should broadly transition together between transcriptionally permissive and repressed states.

Several observations challenge this simple model. First, the genes encoding the core SC are dispersed across independent autosomal loci rather than organised within a common genomic domain, requiring any coordinated regulation to be imposed in trans. Second, transcriptional activation alone cannot ensure assembly of a complex: RNA processing, transcript stability, translation and protein turnover can each reshape the relationship between transcription and final protein abundance. Third, increasing evidence indicates that proteins historically classified as meiosis-specific can be detected outside the germline in cancer (14–19). Somatic expression of individual SC components therefore does not need to represent inappropriate activation of the entire meiotic programme. Instead, it raises the possibility that the apparent meiotic specificity of the SC emerges from the convergence of independently regulated proteins during a restricted developmental window.

Our observations suggest an alternative model in which **SC assembly, rather than SC gene expression, is the coordinated biological event**. We test this model by integrating developmental and stage-resolved germ-cell transcriptomics and proteomics with measurements of nascent transcription, pre-mRNA, mature RNA and protein abundance in somatic cells, together with promoter activity, transcription-factor perturbation, DNA methylation, chromatin accessibility and single-cell transcriptomic analyses across mouse and human tissues. We find that SC genes are neither activated nor extinguished as a synchronous unit during germ-cell development and that RNA and protein abundance become extensively uncoupled across meiotic progression. In somatic cells, individual SC loci occupy distinct transcriptional and chromatin states, while relationships apparent at the level of nascent or pre-mRNA transcription are progressively lost at mature RNA and protein levels. Although DNA methylation can strongly regulate individual SC promoters, endogenous methylation states do not provide a common explanation for SC-gene expression. Finally, across normal mouse and human tissues, individual SC genes show conserved associations with specific somatic cell identities, demonstrating that their expression outside meiosis is structured rather than restricted to aberrant cancer-associated reactivation.

Together, our findings replace a binary model of meiotic ON and somatic OFF states of SC genes with a multilayered regulatory framework in which the components of the SC are independently controlled at transcriptional, RNA and protein levels. We propose that this regulatory uncoupling enables a set of independently encoded proteins to converge transiently to construct a meiosis-specific molecular machine, while permitting individual components to persist or be redeployed in distinct cellular contexts. More broadly, these findings suggest that the developmental specificity of a molecular complex need not arise from coordinated expression of its constituent genes, but can instead emerge from the temporally restricted convergence of independently regulated gene products.

## Results

### Expression of synaptonemal complex genes is regulated by an off-switch in meiosis operating as two modes of regulation rather than one

We first asked whether genes encoding the structural components of the synaptonemal complex behave as a co-regulated transcriptional module. Despite assembling into a single macromolecular structure (**Fig 1a-b**), the eight core SC genes are dispersed across independent autosomal loci (**Fig 1c-d**). Expression of SC genes is turned off prior to meiosis and after its completion apart from *SYCE2* in mice (**Fig 2a-d**). However, analysis of developmental germ-cell transcriptomes revealed that SC activation is not turning on as a single nodule. *SYCP3* and *SYCE1* are the earliest SC genes induced during female germ-cell development, followed by *SYCP2* and subsequently *SYCP1* and the remaining central-element components (**Fig 2a-c**). Similarly in sperm development *SYCP3* is activated first and to the highest extent followed by *SYCP2* and *SYCE1* (**Fig 2d**). Thus, formation of the SC is preceded by ordered activation of independently regulated genes rather than a concerted transcriptional switch. Importantly this gene expression switch is not correlated with protein levels, SC proteins are not absent post SC disassembly and their decrease is not synchronised (**Fig 2e**).

**Figure 1.**
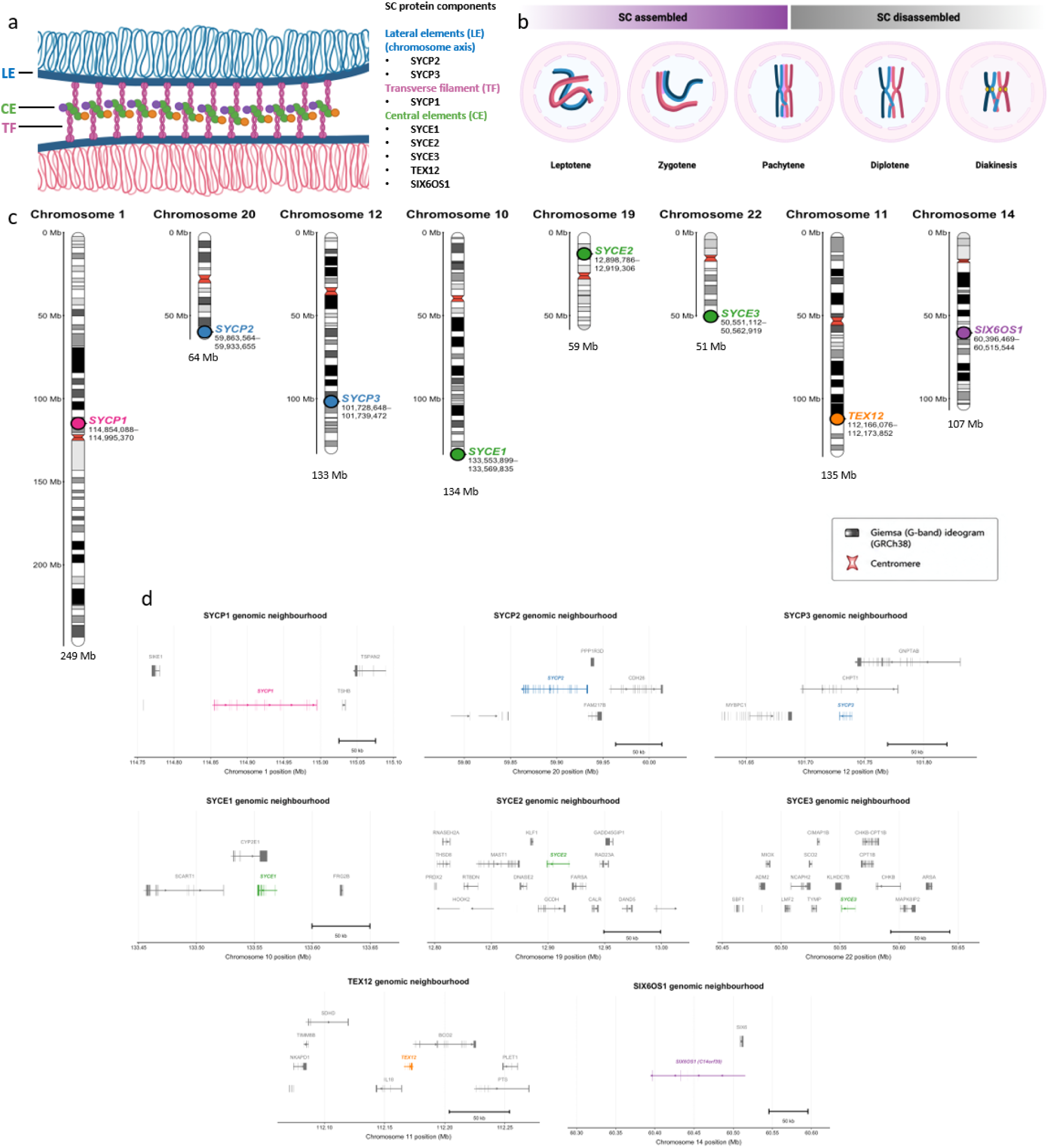
The SC is a physical complex, but its genes are not a genomic module. (**a**) Synaptonemal complex architecture. (**b**) SC dynamics during meiosis. (**c**) Genomic loci of core SC genes in the human genome (GRCh38). (**d**) Genomic environment of synaptonemal complex genes in the human genome (GRCh38).

**Figure 2.**
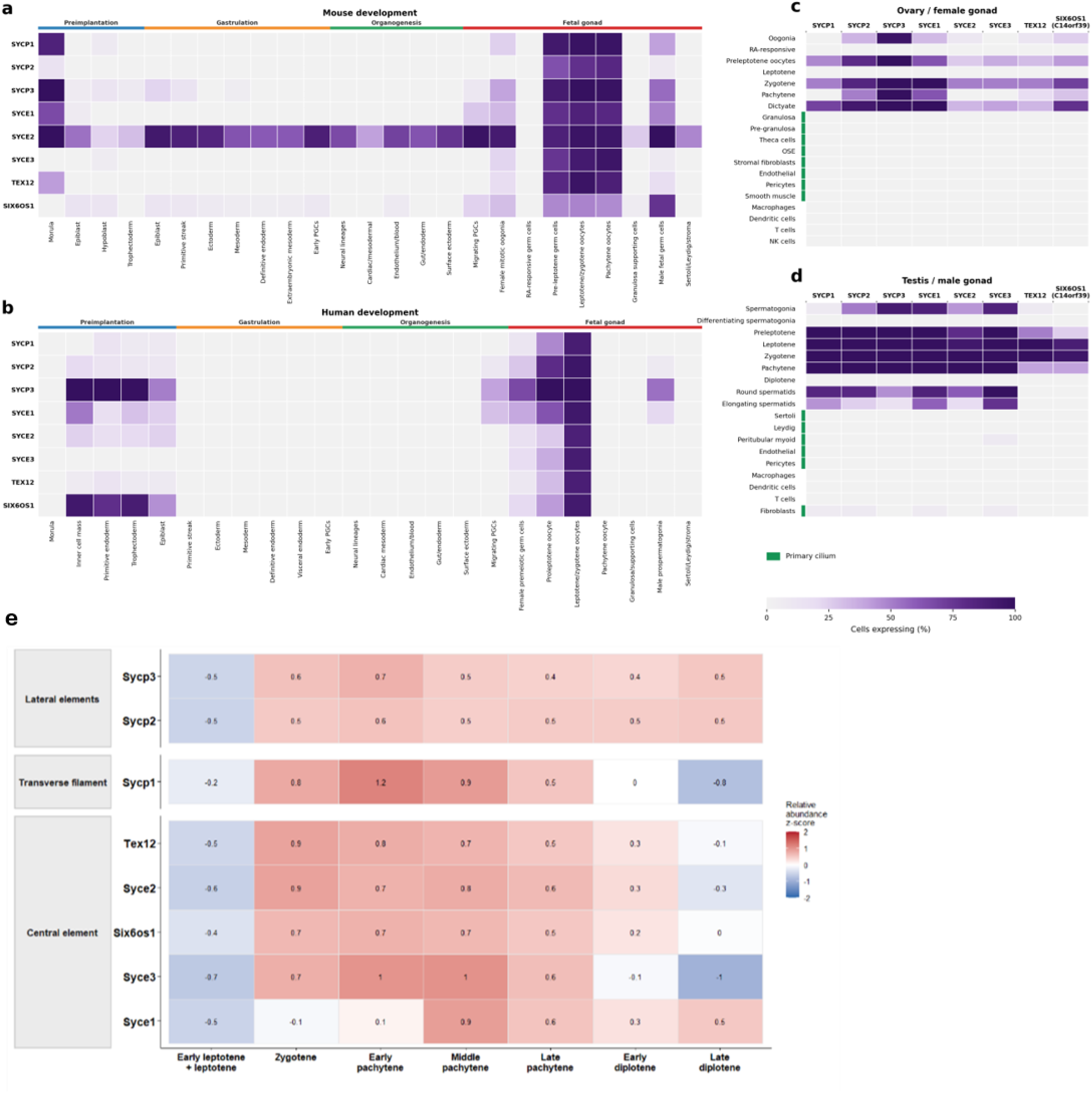
Synaptonemal complex genes are not regulated as a single transcriptional module during meiosis. (**a**) Expression of SC genes during mouse embryonic development. (**b**) Expression of SC genes during human embryonic development. (**c**) Expression of SC genes across human ovarian and female-gonadal cell populations. (**d**) Expression of SC genes across human testicular and male-gonadal cell populations. (**e**) Expression of SC proteins during mouse meiosis.

### Transcript and protein abundance become uncoupled during meiotic progression and SC disassembly

Although levels of most SC proteins decreased following diplotene, individual components persist and SYCP2, SYCP3 and SYCE1 levels increased after diplotene (**Fig 2e**, **3a-b**). Across meiotic stages, RNA levels were not predictive of protein abundance for any SC genes apart from *SYCE1* and to a lesser extent *SYCP1* (**Fig 3c**). Introducing a temporal offset of two stages between RNA and protein measurements improved these relationships (**Fig 3d-f**), consistent with substantial temporal separation between transcription and protein accumulation. This behaviour was not equivalently observed among housekeeping genes (**Fig 3g**) or comparator meiotic gene sets (**Fig 3h**), indicating extensive transcript–protein uncoupling among most SC components.

**Figure 3.**
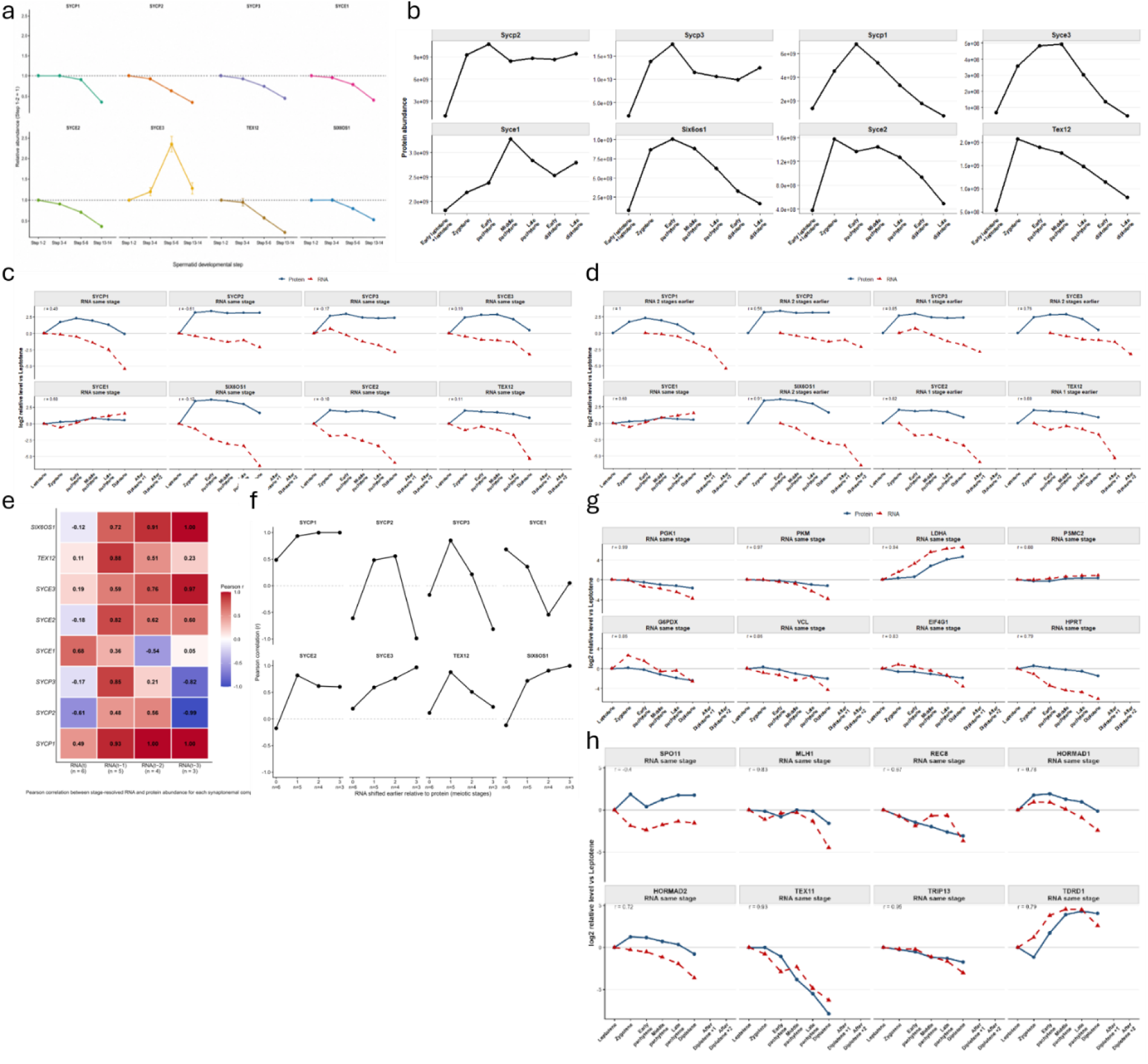
Synaptonemal complex expression is temporally uncoupled during normal germ-cell development. (**a**) SC protein levels in murine spermatids. (**b**) SC protein abundance during murine prophase I. (**c**) SC RNA (blue) and protein (red) levels across murine meiosis. (**d**) SC RNA (blue) and protein (red) levels correlation across murine meiosis after applying a two-stage lag to RNA levels, values are log2 relative to leptotene. (**e-f**) SC RNA-protein level Pearsons correlation heatmap (**e**) and relationship per gene (**f**) across murine meiosis stages with stage matching (t), one-stage (t-1), two stage (t-2) and three stage (t-3) delay applied. (**g-h**) Same stage RNA-protein correlation of house-keeping (**g**) and non-SC meiotic genes (**h**) across murine meiosis.

### Distinct regulatory gates control individual synaptonemal complex genes during meiotic prophase

The discordance between SC transcript and protein abundance suggested that regulation downstream of transcription contributes substantially to the timing of SC protein production. We therefore asked whether the eight SC genes share a common gene regulatory programme or are differentially controlled at distinct steps. We integrated stage-resolved measurements of promoter methylation, chromatin accessibility, active histone modifications, nascent transcription, mature RNA abundance, ribosome engagement and protein abundance across meiotic prophase I (**Fig 4a**). Promoter competence did not constitute a synchronous switch for the SC gene set. Loss of promoter methylation and acquisition of accessible or active chromatin occurred with different kinetics between genes and frequently preceded, or continued after, maximal nascent transcription. *SIX6OS1 (Fig S1)* and *SYCE2 (Fig S3)*, for example, displayed evidence of promoter competence already in leptotene/zygotene, whereas active chromatin marks at several other SC loci were reinforced later in prophase. Thus, promoter demethylation and chromatin accessibility appear to establish permissive states for individual SC loci but do not, by themselves, determine the timing of their transcriptional output.

**Figure 5.**
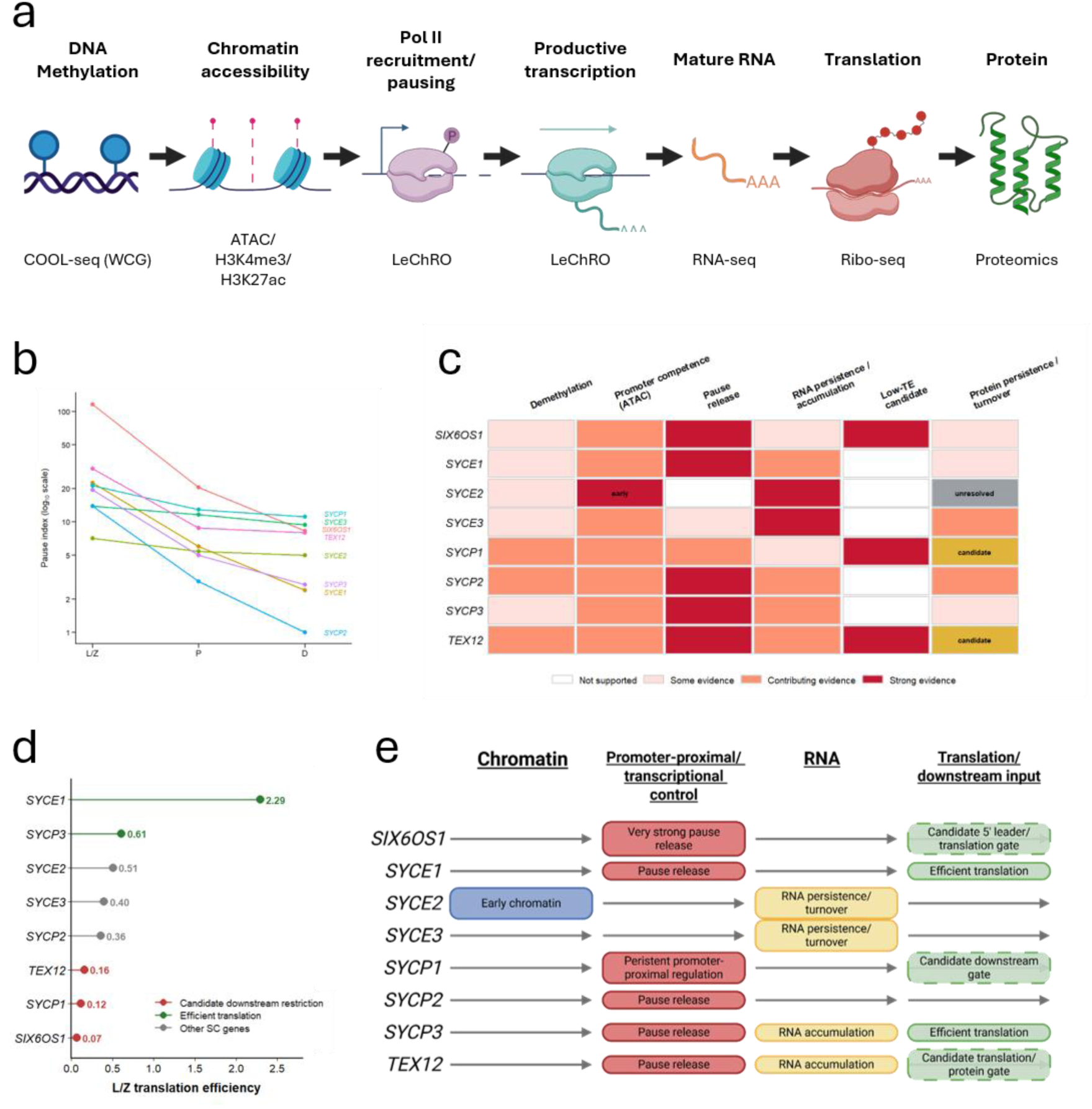
Distinct regulatory gates control individual synaptonemal complex genes during meiotic prophase. **(a)** Regulatory layers interrogated across SC-gene expression, from promoter methylation and chromatin accessibility to transcription, RNA abundance, translation and protein accumulation **(b)** Promoter pause-index trajectories across leptotene/zygotene (L/Z), pachytene (P) and diplotene (D); decreasing values indicate reduced promoter-proximal enrichment relative to gene-body transcription. **(c)** Integrated summary of evidence for regulation at each layer. Colour intensity indicates increasing evidence; candidate and unresolved features are indicated separately. **(d)** Relative L/Z translation efficiency (TE), calculated from matched RNA-seq and Ribo-seq. Low TE identifies SIXcOS1, SYCP1 and TEX12 as candidates for downstream restriction, whereas SYCE1 and SYCP3 show comparatively efficient translation. **(e)** Integrated gene-specific models highlighting the principal regulatory gates identified for each SC gene. Together, these analyses reveal distinct combinations of chromatin, transcriptional, RNA and downstream regulation rather than a single coordinated SC-gene expression programme.

Nascent-transcription profiles revealed a second and unexpectedly heterogeneous regulatory layer. Promoter-proximal enrichment decreased markedly between leptotene/zygotene and pachytene for five SC genes. The pause index decreased from 115.7 to 20.5 for *SIX6OS1*, 22.5 to 6.0 for *SYCE1*, 13.9 to 2.9 for *SYCP2*, 19.4 to 5.0 for *SYCP3* and 30.2 to 8.8 for *TEX12*, with further reductions or stabilization in diplotene (**Fig 4b**). These trajectories are compatible with a transition from promoter-proximal Pol II enrichment into productive elongation during pachynema. In contrast, *SYCE2* showed only a modest change in pause index (7.1 to 5.4 to 5.0), while *SYCE3* similarly retained substantial promoter enrichment (13.9 to 11.6 to 9.4). *SYCP1* displayed an intermediate behaviour, decreasing from 21.2 to 12.9 to 11.1 but retaining substantial promoter-proximal signal. Thus, even within a set of proteins required to assemble the same meiotic structure, productive transcription is not established through an equivalent promoter transition.

Regulatory divergence remained evident downstream of nascent transcription. *SYCP3 (Fig S7)* and *TEX12 (Fig S8)* reached their maximal mature-RNA abundance in diplotene after nascent transcription had peaked in pachytene, consistent with continued RNA accumulation or persistence after maximal transcriptional output. Conversely, *SYCE2 (Fig S3)* and *SYCE3 (Fig S4)* displayed maximal mature and intronic RNA signals already in leptotene/zygotene, preceding their maximal nascent-transcription signal. This reverse temporal relationship cannot be explained by a simple transcription-to-mRNA processing delay and instead implicates differences in pre-existing RNA pools, RNA persistence or turnover, and/or developmental population history. *SIX6OS1 (Fig S1)*, *SYCE1* (*Fig S2*) and *SYCP2 (Fig S6)* similarly retained substantial RNA after nascent transcription declined. Collectively, these patterns demonstrate that mature SC transcript abundance cannot be inferred directly from contemporaneous transcriptional activity.

Translation provided a further point of divergence. Matched RNA-seq and Ribo-seq in leptotene/zygotene revealed more than a 30-fold range in relative translation efficiency among SC transcripts. *SYCE1* showed the highest translation efficiency of the eight genes (TE = 2.29), followed by *SYCP3* (0.61), whereas *TEX12* (0.16), *SYCP1* (0.12) and *SIX6OS1* (0.07) occupied the lowest positions in the SC-gene ranking (**Fig 4c**). High relative translation efficiency for *SYCE1* and *SYCP3* argues against inefficient translation constituting their principal restrictive gate. In contrast, low translation efficiency identifies *SIX6OS1*, *SYCP1* and *TEX12* as candidates for additional downstream regulation. For *SIX6OS1*, this interpretation is further supported by ribosome accumulation within the 5′ leader, although the present data do not distinguish productive upstream translation from initiation, pausing or ribosome stalling (*Fig S1*).

Integration of these regulatory layers revealed that no single step in the expression cascade accounts for the expression of all eight SC genes (**Fig 4d,e**). *SIX6OS1, SYCE1, SYCP2, SYCP3* and *TEX12* show the strongest evidence for a promoter-proximal transition into productive transcription, whereas RNA persistence or turnover is particularly prominent for *SYCE2* and *SYCE3. SYCP3* and *TEX12* additionally show later mature-RNA accumulation, while *SIX6OS1, SYCP1* and *TEX12* emerge as the strongest candidates for downstream translational or protein-turnover control. Importantly, DNA demethylation, chromatin accessibility, transcription, RNA accumulation and translation do not define successive synchronous states of a single SC expression programme.

Together, these results indicate that assembly of the synaptonemal complex is achieved by convergence of gene products where expression is controlled through different combinations of regulatory mechanisms. Rather than being switched on as a transcriptionally unified module, individual SC genes acquire promoter competence, productive transcription, stable RNA and translational output through distinct trajectories. This multilayered regulatory architecture provides a mechanistic explanation for the transcript–protein uncoupling observed during meiotic progression and establishes that coordinated assembly of the SC does not require coordinated regulation of its constituent genes.

### Synaptonemal complex genes occupy distinct transcriptional states in somatic cells

Having established that SC genes are independently regulated across meiotic development, we next asked whether their expression becomes coordinated when they are re-expressed in somatic cells. We detected expression of all 8 SC proteins in MCF7 cells (*Fig S9*) making them a good model for comparison. We analysed phase-resolved nascent transcription in MCF7 cells together with promoter accessibility, active chromatin marks and RNA polymerase II occupancy. Nascent transcription was detectable for seven of the eight SC genes, demonstrating that expression of multiple SC components in cancer cells reflects active transcription rather than exclusively persistence of pre-existing RNA (**Fig 5**). However, both the distribution and cell-cycle dynamics of engaged transcription differed markedly between individual SC loci (**Fig 5**).

**Figure 5.**
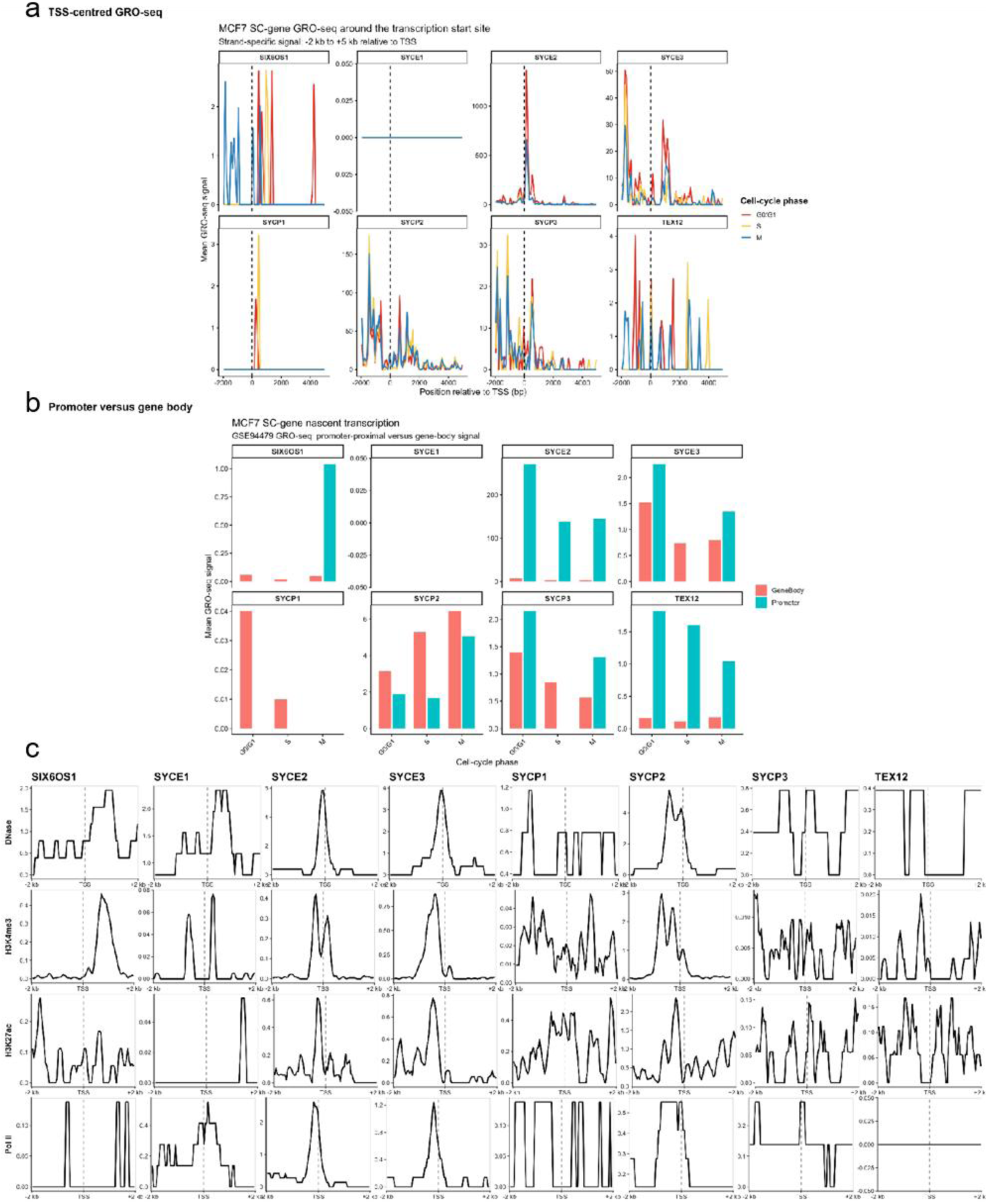
Phase-resolved nascent transcription and promoter chromatin reveal heterogeneous regulatory states among SC genes in somatic cells. **(a)** TSS-centred strand-specific GRO-seq profiles for the eight SC genes across G0/G1, S and G2/M phases in MCF7 cells. **(b)** Mean promoter-proximal and gene-body GRO-seq signal across the same phases in MCF7 cells. GRO-seq data were mined from GEO GSES447S, with two replicates averaged per phase. **(c)** Bulk MCF7 promoter-centred DNase, H3K4me3, H3K27ac and RNA polymerase II profiles across ±2 kb of the selected TSS.

*SYCE2* displayed the clearest conventional transcriptionally active state. A strong promoter-proximal GRO-seq peak coincided with DNase accessibility, H3K4me3 and Pol II occupancy at the same transcription start site, providing concordant evidence of promoter recognition and transcriptional engagement (**Fig 5a-c**). *SYCP2* was similarly associated with an accessible, H3K4me3-positive, Pol-II-occupied promoter region, but showed a fundamentally different transcriptional profile: nascent RNA extended broadly across the gene body and increased towards G2/M. *SYCE3* also showed concordance between nascent transcription and a chromatin-supported canonical promoter, although its phase-resolved transcriptional profile was more complex (**Fig 5a-c**). Thus, even among SC genes with strong evidence of transcriptionally competent promoters, transcription was organised differently.

Other SC loci occupied states in which promoter competence and productive transcription were strikingly uncoupled. *SYCE1* represented the clearest example. Despite detectable DNase accessibility, H3K4me3 and Pol II occupancy at a candidate alternative promoter, no productive nascent transcription was detected in the final GRO-seq profiles (**Fig 5a-c**). This configuration is compatible with a promoter that can be accessed and recruit transcriptional machinery but does not progress to substantial productive transcription. Conversely, *TEX12* showed the opposite discordance: promoter-associated nascent transcription was reproducibly detected despite the absence of official DNase, H3K4me3, H3K27ac or Pol II peaks at the retained reference promoter in the bulk MCF7 chromatin datasets (**Fig 5a-c**). *SIX6OS1* provided a third regulatory configuration, in which sparse nascent transcription was accompanied by stronger promoter-like chromatin at an alternative transcription start site than its canonical start (**Fig 5a-c**). Thus, SC genes expressed within the same somatic cell population occupy multiple distinct regulatory states rather than sharing a common reactivation mechanism.

### SC gene expression is progressively uncoupled across transcriptional, RNA and protein regulatory layers

We quantified pre-mRNA, mature mRNA and protein abundance for the eight major SC genes, *SYCP1, SYCP2, SYCP3, SYCE1, SYCE2, SYCE3, TEX12 and SIX6OS1*, across a diverse panel of human cell lines. Pre-mRNA and mature mRNA were measured independently by qRT-PCR and expressed as ΔCt values, while protein abundance was quantified from immunoblotting. For comparisons between regulatory layers, RNA measurements were converted to relative abundance (2^−ΔCt), such that increasing values represented increasing molecular abundance across all three datasets.

Analysis of pre-mRNA abundance revealed that SC genes are not transcribed as one regulatory unit. Pairwise correlations identified substantial co-expression between specific subsets of SC genes including correlation between *SYCP2, SYCE2*, and *SYCE3* as well as *TEX12* and *SIX6OS1* (r=0.94) (**Fig 6a**). Hierarchical clustering consequently resolved partially overlapping transcriptional modules, including a *SYCP2–SYCE2–SYCE3* group and a *SYCE1–TEX12–SIX6OS1* group with *SYCP3* particularly associated with *SIX6OS1*. *SYCP1* is the most weakly coupled member overall and appears comparatively independently regulated. Thus, despite residing at independent genomic loci, subsets of SC genes can share transcriptional behaviour across somatic cellular backgrounds. Importantly, however, the eight genes did not form a single transcriptionally coordinated unit.

**Figure 6.**
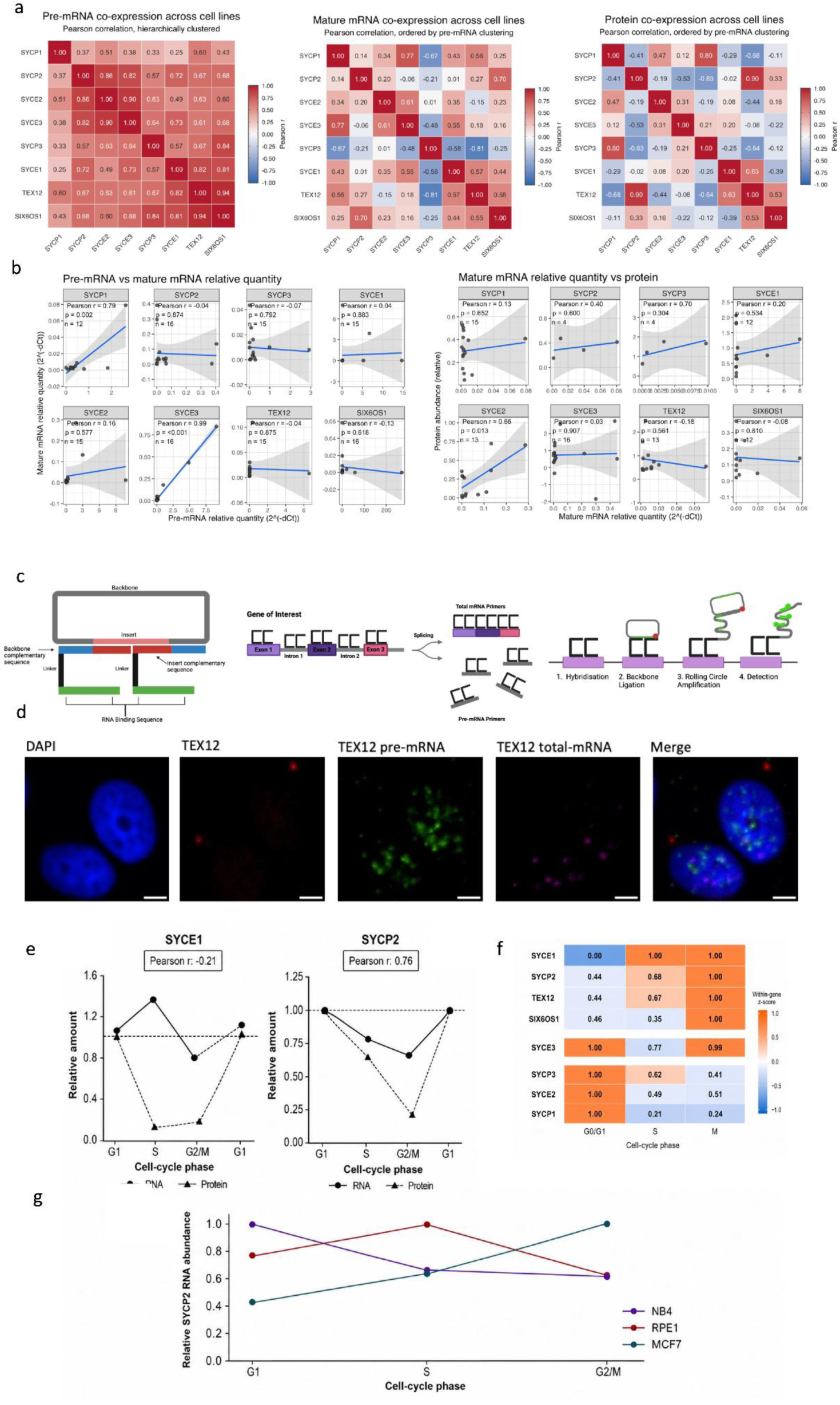
SC protein levels are regulated post transcriptionally in somatic cells. (**a**) Pearson correlation heatmaps showing co-expression between SC genes at the pre-mRNA, mature mRNA and protein levels across somatic cell lines tested. Pre-mRNA expression was hierarchically clustered, with the same gene ordering applied to mature mRNA and protein datasets. (**b**) Correlation of pre-mRNA to mature mRNA expression and mature mRNA to protein abundance across individual SC genes in somatic cell lines. Pearson correlation coefficients, p-values and number of paired cell lines (n) are shown for each gene; shaded areas indicate S5% confidence intervals. (**c**) Schematic overview of PLAYR probe design, targeting of pre-mRNA and total mRNA, and rolling-circle amplification for fluorescent RNA detection. (**d**) Representative PLAYR images of MCF7 cells showing TEX12 protein at centrosomes together with TEX12 pre-mRNA and total mRNA (**e**) NB4 RNA and protein abundance of SYCE1 and SYCE2 across cell cycle. (**f**) Relative SC gene GRO-seq transcription across the MCF7 cell cycle. (**g**) Relative SYCP2 RNA abundance during NB4, RPE1 and MCF7 cell cycles.

This organisation was substantially altered when mature mRNA rather than pre-mRNA was examined. Correlations between SC genes were generally weaker and the relationships identified at the pre-mRNA level were reorganised (**Fig 6a**). Most notably, *SYCP3* mature RNA abundance was negatively correlated with several other SC transcripts, while other associations present among pre-mRNAs were reduced or lost. Therefore, the relative abundance of nascent SC transcripts does not simply propagate into the mature RNA pool. These observations suggested that transcript processing and/or stability represents an important additional regulatory layer that can reshape the transcriptional relationships between SC genes.

Protein abundance introduced a further level of regulatory divergence. Hierarchical clustering of protein measurements generated a pattern distinct from either RNA layer, demonstrating that transcriptional co-expression does not necessarily predict coordinated protein accumulation (**Fig 6a**). We therefore directly compared pre-mRNA and mature mRNA abundance for individual genes (**Fig 6b**). Surprisingly, for most SC genes these two measurements were poorly correlated. *SYCP2, SYCP3, SYCE1, SYCE2, TEX12* and *SIX6OS1* showed little or no linear relationship between pre-mRNA and mature mRNA abundance across the analysed cell lines. For example, *SYCE1* (r = 0.04), *SYCE2* (r = 0.16), *TEX12* (r = −0.04) and *SIX6OS1* (r = −0.13) showed essentially no correspondence between the two RNA pools. *SYCE3* was a striking exception, displaying an almost complete correspondence between pre-mRNA and mature mRNA abundance (r = 0.99, p < 0.0001). Thus, coupling between transcriptional output and mature transcript accumulation is itself gene-specific within the SC genes. Mature mRNA–protein comparisons reinforced this conclusion but also revealed marked gene-specific differences (**Fig 6b**). Mature *SYCE1* mRNA showed little relationship with protein abundance (r = 0.20). *TEX12* showed a similar pattern with no relationship between mature mRNA and protein (r = −0.18). For both genes, therefore, steady-state mature mRNA abundance was not predictive of protein accumulation. *SYCE2* showed a different regulatory architecture. Although its pre-mRNA and mature mRNA levels were poorly correlated (r = 0.16), protein abundance correlated with mature mRNA (r = 0.66, p = 0.014). Conversely, *SYCE3*, despite showing exceptionally tight coupling between pre-mRNA and mature mRNA, displayed a different relationship at the protein level. Mature mRNA did not correlate with protein abundance (r = 0.03). Thus, even near-perfect transmission of transcriptional output into the mature RNA pool does not necessarily result in proportional protein accumulation.

Other SC genes showed further uncoupling. *SIX6OS1* displayed no detectable relationship between pre-mRNA and mature mRNA (r = −0.13), pre-mRNA and protein (r = −0.05), or mature mRNA and protein (r = −0.08). *SYCP2* similarly showed no relationship between its pre-mRNA and mature transcript abundance (r = −0.04). *SYCP3* also showed essentially no relationship between pre-mRNA and mature mRNA (r = −0.07). Together, these data reveal that SC gene expression is progressively remodelled across transcriptional, RNA and protein layers. Genes encoding proteins that ultimately assemble into the same macromolecular structure can be transcriptionally co-regulated, yet these relationships can be weakened, lost or reorganised at the level of mature RNA and reorganised again at the protein level. Moreover, individual SC genes follow distinct regulatory trajectories: *SYCE3* maintains exceptionally tight pre-mRNA–mRNA coupling, *SYCE2* retains relationships between both RNA pools and protein, whereas *SIX6OS1* and *SYCP2-3 are* extensively uncoupled across all three measured layers. We confirmed expression of all regulatory levels by PLAYR proximity ligation for RNA assay (20, 21) modified to cell imaging with protein colocalization (**Fig 6c-d**). These findings identify post-transcriptional regulation as a major determinant of SC protein expression and provide a mechanistic framework through which individual components of a normally coordinated meiotic structure could be selectively retained or re-expressed outside meiosis without activation of the complete SC programme. Similar discordance was observed following cell-cycle fractionation and across distinct somatic cell backgrounds (**Fig 6e-g**), demonstrating that transcriptional co-regulation is neither sufficient nor necessary to generate coordinated SC protein abundance.

### DNA methylation regulates individual SC promoters but does not constitute a common somatic silencing mechanism

Reporter analysis demonstrated that multiple SC promoters are transcriptionally competent in somatic cells and can respond to retinoic-acid signalling (**Fig 7a-b**). Our analysis showed that of analysed SC genes only *SYCP3* was significantly methylated in somatic cells and was regulated by methylation (**Fig 7c, h, l, n**). Experimental promoter methylation was also able to suppress *SYCE3* and *SYCE2* promoters and could be reversed by pharmacological inhibition of DNA methylation (**Fig 7d-e**). However, endogenous promoter methylation varied extensively between SC genes and cell lines and failed to predict transcript or protein abundance (**Fig 7f-m**). Together with chromatin-accessibility (**Fig 7n**) and nascent-transcription measurements (**Fig 5**), these observations demonstrate that promoter methylation can regulate individual SC loci but cannot explain the coordinated restriction or somatic re-expression of the SC gene family.

**Figure 7.**
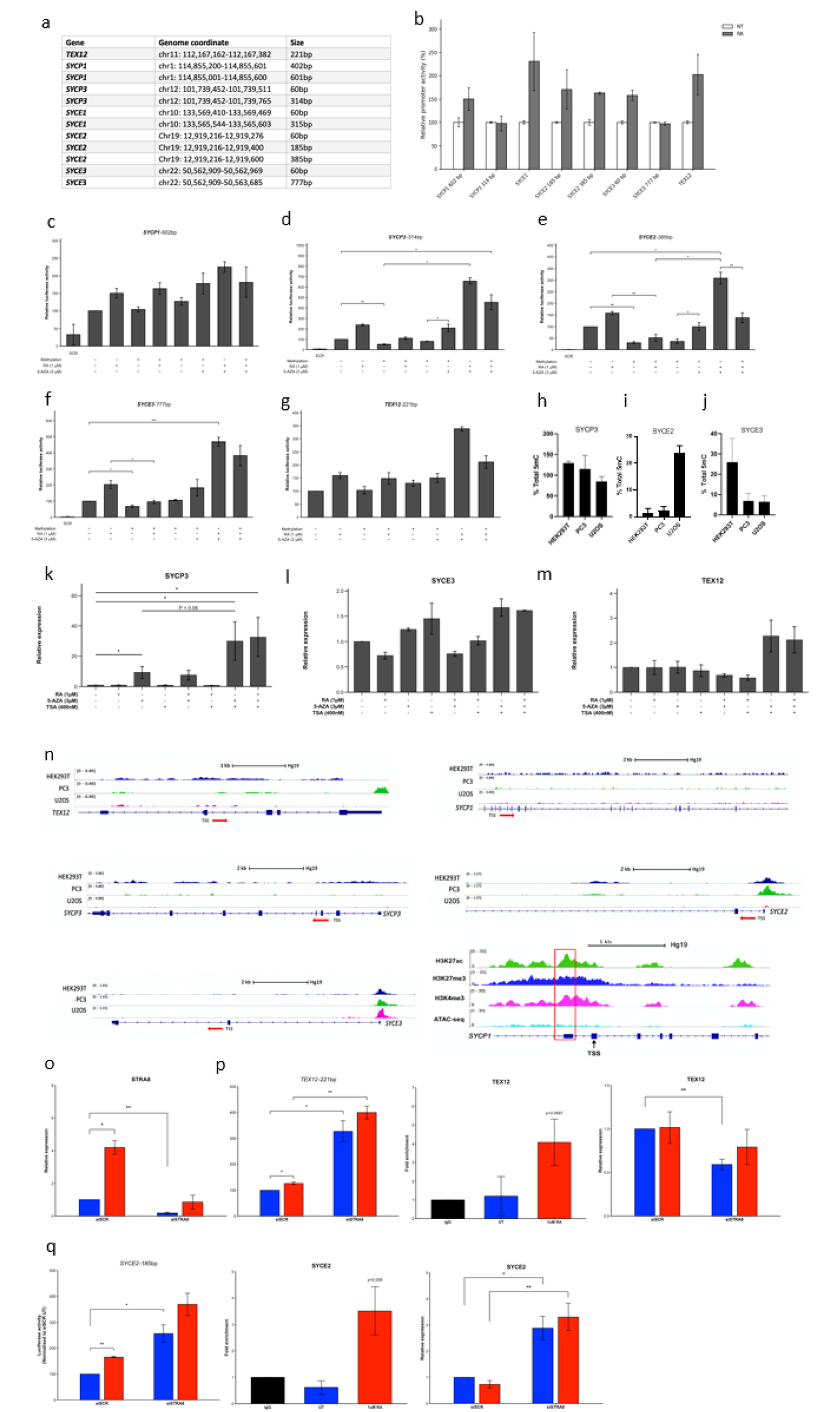
Promoter methylation does not explain endogenous SC expression. (**a**) Identified SC genes promoter regions. (**b**) Luciferase reporter assay in HEK2S3 cells showing SC promoter activity in luciferase reporter assay as well as response to RA stimulation. (**c-g**) HEK2S3 cells transfected with SC promoter driven luciferase construct with and without prior methylation as indicated. (**h-j**) Cell’s DNA was extracted and 5mC methylation levels were detected by Gluc-PCR (22). (**k-m**) HEK2S3 cells were treated as indicated and following RNA extraction SC transcript levels were measured with qRT-PCR normalised to HPRT1. (**n**) ATAC-seq analysis of SC gene chromatin accessibility in somatic cells. (**o**) STRA8 transcript levels following RA treatment and STRA8-targeting siRNA. (**p**) Luciferase reporter assay of the TEX12 promoter following STRA8 silencing, STRA8 ChIP-qPCR following RA stimulation, and TEX12 transcript levels following STRA8 knockdown. (**q)** Luciferase reporter assay of the SYCE2 promoter following STRA8 silencing, STRA8 ChIP-qPCR following RA stimulation, and SYCE2 transcript levels following STRA8 knockdown

We next asked whether SC promoters that respond to retinoic acid (RA) signalling are directly regulated by the meiotic transcriptional regulator STRA8, and whether promoter regulation is propagated to endogenous transcript abundance. In HEK293 cells, silencing of *STRA8* increased *TEX12* promoter activity in luciferase reporter assays, indicating that STRA8 negatively influences the activity of the isolated *TEX12* promoter (**Fig 7o-p**). Consistent with direct promoter engagement, ChIP–qPCR detected STRA8 recruitment to the endogenous *TEX12* promoter following RA treatment, whereas promoter occupancy was not detected in untreated cells (**Fig 7p**). Thus, RA induces recruitment of STRA8 to the *TEX12* promoter, and perturbation of *STRA8* is sufficient to alter its transcriptional activity in a promoter-reporter context.

However, this relationship was not reproduced at endogenous *TEX12* locus. STRA8 silencing produced a modest reduction in *TEX12* transcript but this difference did not reach statistical significance (**Fig 7p**). The direction of this response was therefore opposite to that observed using the isolated promoter reporter. These results indicate that STRA8 occupancy and promoter responsiveness alone are insufficient to predict steady-state *TEX12* transcript abundance, consistent with additional regulation operating within the endogenous genomic and/or post-transcriptional context.

*SYCE2* showed a superficially similar promoter response but a distinct endogenous transcriptional outcome. As observed for *TEX12*, *STRA8* silencing increased *SYCE2* promoter activity in luciferase assays, and ChIP–qPCR demonstrated recruitment of STRA8 to the endogenous *SYCE2* promoter following RA treatment (**Fig 7q**). In contrast to *TEX12*, however, *STRA8* silencing resulted in a significant increase in endogenous *SYCE2* transcript abundance. Thus, for *SYCE2*, the effect of *STRA8* depletion on the isolated promoter was retained at the level of steady-state endogenous RNA, consistent with STRA8 contributing directly or indirectly to repression of *SYCE2* expression in this cellular context (**Fig 7q**).

### SC genes are selectively and reproducibly redeployed in mammalian somatic cell types

Finally, analysis of single-cell transcriptomes across human and mouse tissues revealed that somatic SC expression is neither restricted to cancer nor explained by stochastic leakage of the meiotic programme (**Fig 8)**. Individual SC genes showed conserved associations with specific somatic cell identities across organs and species, including *SYCE1* expression in fibroblasts (**Fig 8a**), *SYCE3* in dendritic (**Fig 8b**) and macrophage (**Fig 8c**) populations. These conserved patterns demonstrate that components of the meiotic SC programme can be independently incorporated into somatic transcriptional programmes.

**Figure 8.**
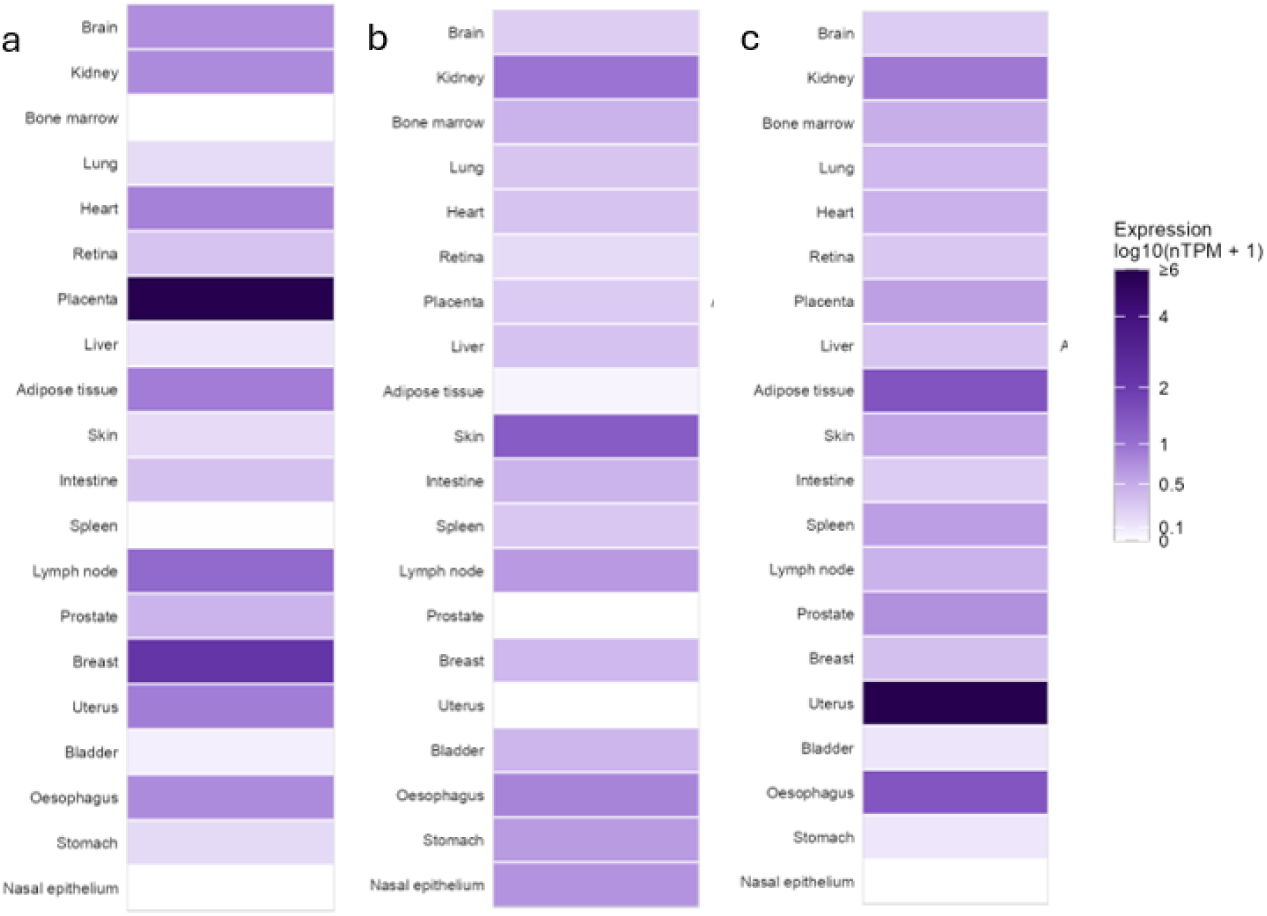
Individual SC genes are reproducibly redeployed in specific somatic cell identities. (**a**) SYCE1 transcript levels across fibroblasts from sc-RNAseq of 26 organs in human. (**b**) SYCE3 transcript levels across dendritic cells from sc-RNAseq of 26 organs in human. (**c**) SYCE3 transcript levels across macrophages from sc-RNAseq of 26 organs in human.

### Discussion - Multilayered regulatory uncoupling enables modular deployment of synaptonemal complex proteins

The synaptonemal complex is one of the most specialised molecular structures assembled by mammalian cells. Its eight core components converge during meiotic prophase to form a highly ordered chromosome-spanning structure, and their disappearance as cells exit meiotic prophase has encouraged an intuitive model in which SC genes constitute a coordinated meiotic programme: switched on during meiosis, switched off thereafter, and maintained in an epigenetically repressed state in somatic cells (1, 5, 6, 8). This view parallels the broader cancer/testis gene paradigm, in which germline genes have frequently been considered transcriptionally silent in normal soma and aberrantly reactivated in cancer, with promoter DNA methylation proposed as a major determinant of this switch (11, 13, 23–25). Our results highlight that this framework is insufficient for the SC. Rather than behaving as a single regulatory unit, SC genes are independently controlled at multiple stages between promoter competence and protein accumulation. Their defining feature is therefore not synchronous gene expression, but the transient convergence of independently regulated gene products to construct the SC. The eight mammalian SC genes are distributed across independent autosomal loci and encode proteins occupying distinct positions within the SC: SYCP2 and SYCP3 form lateral-element components, SYCP1 forms transverse filaments, and SYCE1, SYCE2, SYCE3, TEX12 and SIX6OS1 contribute to the central element (**Fig 1**). Their genomic dispersion does not preclude coordinated regulation by trans-acting mechanisms, but it requires such coordination to be imposed independently at multiple loci. Our data show that this does not occur through a simple simultaneous switch. SC transcripts appear sequentially during meiotic entry, with different components becoming detectable at different developmental stages (**Fig 2**). Likewise, their expression does not terminate synchronously when the mature SC is no longer required (**Fig 2e**, **3**). Thus, even within the physiological context for which these proteins are best understood, SC expression is not a single molecular state.

The separation becomes apparent when transcription and protein abundance are considered independently. SC transcripts persist beyond the period of SC assembly (**Fig 2**), while individual proteins follow markedly different trajectories following pachytene and diplotene (**Fig 3**). For most SC genes, RNA abundance does not predict protein abundance in meiosis (**Fig 3**). Importantly, this behaviour is not reproduced by housekeeping genes, broader meiotic gene sets or comparator components of other meiotic structures (**Fig 3g-h**). The disconnect therefore cannot be explained simply by the general temporal complexity of meiotic differentiation. Instead, it suggests that SC biogenesis relies extensively on temporal separation between transcription, RNA availability and final protein accumulation.

Our analysis provides a mechanistic basis for this uncoupling. Promoter demethylation, accessibility and active histone modifications occur with different kinetics between SC loci and frequently precede, or follow, nascent transcription (**Fig 4**). Moreover, five genes, *SIX6OS1, SYCE1, SYCP2, SYCP3* and *TEX12*, show strong decreases in promoter-proximal enrichment as cells progress into pachynema, compatible with release into productive elongation, whereas *SYCE2* and *SYCE3* have much weaker transitions (*Fig S1-8*). These observations are consistent with reports that promoter-proximal RNA polymerase II regulation is an important component of the meiotic transcriptional programme (26, 27), but reveal that individual SC loci participate in this programme to very different extents. Promoter competence and productive transcription are separable regulatory events for SC genes.

Regulatory divergence continues downstream of transcription. *SYCP3* and *TEX12* acquire mature-RNA later following their pachytene nascent-transcription peaks, whereas *SYCE2* and *SYCE3* display substantial mature-RNA pools before maximal nascent transcription (**Fig 4**). The latter behaviour cannot be explained by a simple forward delay from transcription through processing and instead implicates RNA persistence or turnover. Translation introduces another layer of heterogeneity. Among the eight genes, matched leptotene/zygotene translation efficiencies span more than 30-fold, from high relative efficiency for *SYCE1* to very low values for *SIX6OS1, SYCP1* and *TEX12* (**Fig 4d**). Ribo-seq, together with RNA–protein divergence, identify translation and/or protein turnover as plausible additional regulatory gates for selected SC components. The emerging picture is therefore not a linear cascade in which chromatin opening produces RNA and RNA predicts protein, but a series of gene-specific regulatory filters that repeatedly reshape SC output.

This architecture may help explain a long-standing paradox surrounding germ-cell genes in cancer. Cancer/testis genes have historically been interpreted predominantly through the lens of epigenetic derepression (11, 13, 23–25). Promoter hypomethylation is unquestionably important for many classical cancer/testis genes, particularly members of the MAGE family, and pharmacological demethylation can induce their expression (28, 29). However, large-scale analyses have also shown that tumours generally express only subsets of germline genes rather than reinstating a complete germ-cell programme (30–32). Our findings provide a potential explanation for this selectivity. If SC components are independently regulated even in germ cells, there is no requirement for their somatic re-expression to occur as a unit.

Indeed, our somatic-cell analyses reveal several fundamentally different regulatory states among SC genes within the same cellular background. In MCF7 cells, *SYCE2*, *SYCE3* and *SYCP2* show relatively conventional combinations of nascent transcription and promoter-associated chromatin, whereas *SYCE1* provides the contrasting example of promoter-like accessibility, H3K4me3 and Pol II occupancy without detectable productive transcription (**Fig 5**). *SIX6OS1* shows evidence consistent with altered promoter usage, while *SYCP1* and *TEX12* exhibit nascent-transcription features without equivalently strong bulk promoter signatures (**Fig 5**). Thus, even within one cancer-cell population there is no identifiable “SC-gene chromatin state”. Instead, individual loci can be accessible, poised, productively transcribed, alternatively initiated or weakly and conditionally active. The same conclusion emerges from our measurements across successive expression layers. Relationships between subsets of SC genes can be detected at the pre-mRNA level, indicating that some loci share transcriptional inputs (**Fig 6a**). Yet these relationships largely disappear at the mature-RNA level and do not predict protein abundance (**Fig 6a-b**). Moreover, cell-cycle-associated transcriptional behaviour is context dependent: SC genes that resemble proliferation-associated expression patterns in one cell line do not do so in another (**Fig 6e-g**). This suggests that individual SC loci can be incorporated into different regulatory programmes depending on cellular context.

Our promoter experiments reinforce this distinction. Experimental methylation can inhibit several SC promoters (**Fig 7c-m**), demonstrating that DNA methylation is capable of regulating their transcriptional competence. However, endogenous methylation does not consistently predict expression across SC genes or somatic cell lines (**Fig 7**). Some expressed loci are largely unmethylated, others retain appreciable promoter methylation, and chromatin accessibility, nascent transcription, mature RNA and protein abundance frequently fail to align. DNA methylation should therefore be considered one regulatory input rather than the master switch defining SC identity. For SC genes, our data support combinatorial regulation involving promoter identity, transcription-factor recruitment, chromatin state, productive elongation, RNA persistence, translation and protein turnover.

The responses to retinoic-acid signalling provide an informative example of this multilayer organisation. SC promoter reporters are transcriptionally competent in somatic cells and respond differentially to perturbation of retinoic-acid receptors and STRA8 (**Fig 7**). Yet changes in promoter activity do not invariably propagate to equivalent changes in endogenous mature RNA. Conversely, for *SYCE2*, promoter regulation is accompanied by changes in endogenous transcript abundance (**Fig 7q**). The presence of STRA8 at responsive promoters therefore does not imply that all downstream regulatory barriers have been removed. These observations highlight a general limitation of inferring gene expression from promoter activity alone: two loci can receive related transcriptional inputs yet produce different RNA and protein outputs because regulatory information is subsequently modified at downstream layers.

The biological relevance of this independence becomes apparent in normal somatic tissues. Germ-cell proteins have historically been classified as tissue restricted using approaches such as bulk tissue RNA measurements and immunoblotting (1, 5, 6, 8). Such experiments were appropriate for establishing strong tissue enrichment but have an intrinsic limitation: an organ containing millions of cells can contain a biologically important population expressing a gene without producing sufficient bulk signal for detection. A protein expressed reproducibly in a rare epithelial, stromal, neuronal or immune-cell population can therefore appear “absent” when the tissue is treated as a homogeneous sample. Single-cell transcriptomics changes the resolution at which meiotic specificity can be tested. Rather than asking whether an SC transcript is detectable in the whole brain, heart, lung or intestine, it becomes possible to ask whether it is reproducibly expressed by a particular cellular identity across those organs. Our cross-tissue analysis reveals that individual SC genes show recurring associations with specific somatic populations across human tissues, including *SYCE1* with fibroblast populations and *SYCE3* with dendritic cells and macrophages (**Fig 8**). These patterns are difficult to reconcile with random transcriptional leakage. Their recurrence across anatomical sites suggests that at least some SC genes might participate in conserved somatic cellular programmes.

This observation raises the possibility that the historical category of “meiosis-specific protein” contains two biologically distinct concepts: proteins whose expression is genuinely restricted to meiotic cells and proteins/complexes whose meiotic function is restricted to meiosis but whose individual components can be deployed elsewhere. SC proteins have been considered representatives of the former category. Increasing functional evidence suggests that at least some belong to the latter. TEX12 can localise to centrosomes independently of chromosome synapsis and its somatic expression influences centrosome biology, demonstrating that an SC component can acquire a cellular context fundamentally different from its meiotic chromosome function (18). More broadly, aberrantly expressed meiotic proteins can influence DNA repair, chromosome segregation and genome stability in cancer (16, 17, 33–35). Such observations argue that somatic expression should not automatically be interpreted as biologically inert “leakiness”.

Our findings also suggest that somatic moonlighting may not be exclusively a pathological consequence of cancer. If specific SC genes are reproducibly retained or reactivated in defined normal cell types, some functions currently interpreted as cancer-specific repurposing may derive from physiological capacities already used by specialised somatic cells. Together, these findings support a model in which SC genes are not controlled by a binary meiotic ON/somatic OFF switch. Their expression is independently gated through promoter selection and chromatin state, transcription, RNA maturation or stability, translation and protein persistence. This architecture permits eight independently regulated gene products to converge transiently during meiosis to construct the synaptonemal complex, while subsequently allowing individual components to be retained, extinguished or redeployed into distinct somatic cellular programmes.

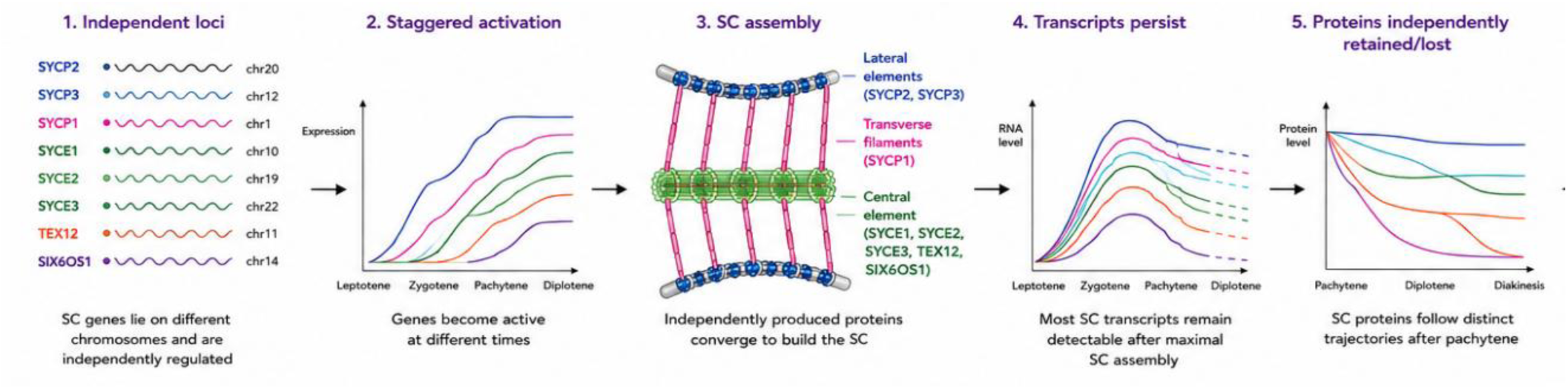

## Materials and methods

### Cell culture

HEK293T, U2OS, MCF7, A375, AGS, MIA-Pa-Ca-2, DU145, HeLa, JIMT-1, WM1361, WM164, U251 and COS-7 cells were maintained in DMEM supplemented with 10% (v/v) foetal bovine serum (FBS). BxPC-3, LNCaP, PC3, CWR22Rv1, THP-1, Omm2.3, COR-L23, PC9, U937, PEO1, PEO4, Mel270, AsPC-1 and NALM-6 cells were maintained in RPMI 1640 supplemented with 10% FBS. MV-4-11 and MOLM-13 cells were maintained in RPMI 1640 supplemented with 20% FBS. RPE1 cells were maintained in a 1:1 mixture of DMEM and Ham’s F-12 medium supplemented with 10% FBS. T84 cells were maintained in Ham’s F-12 medium supplemented with 10% FBS. HepG2, Hep3B and SKMEL1 cells were maintained in EMEM supplemented with 10% FBS, whereas Caco-2 cells were maintained in EMEM supplemented with 20% FBS. HL-60 cells were maintained in IMDM supplemented with 20% FBS. HT29 cells were maintained in modified McCoy’s 5A medium supplemented with 10% FBS, and LoVo cells were maintained in F-12K medium supplemented with 10% FBS. MCF10A cells were maintained in MEGM BulletKit medium (Lonza, CC-3150) supplemented with 100 ng/mL cholera toxin.

All cells were cultured in the presence of 1% PenStrep. Cells were maintained at 37°C in a humidified incubator containing 5% CO₂ and passaged at an appropriate confluency using trypsin–EDTA.

### Western blotting

Protein samples were prepared in 2× Laemmli Sample Buffer (Bio-Rad, cat. no. 1610737) supplemented with 10% (v/v) β-mercaptoethanol and boiled at 100°C for 10 min. Proteins were resolved by SDS–polyacrylamide gel electrophoresis using 10–15% resolving gels and transferred onto nitrocellulose membranes in transfer buffer containing 10% (v/v) methanol. Membranes were blocked in 5% (w/v) milk and incubated with the indicated primary antibodies, followed by the appropriate HRP-conjugated secondary antibodies. Protein bands were detected using Clarity Max Western ECL Substrate (Bio-Rad, cat. no. 1705062) and chemiluminescent signals were acquired using a ChemiDoc Imaging System (Bio-Rad).

### Reverse transcription and qPCR

Total RNA was extracted from cultured cells using the RNeasy Plus Mini Kit (Qiagen, cat. no. 74134) in combination with QIAshredder columns (Qiagen, cat. no. 79654), according to the manufacturer’s instructions. Samples were subjected to on-column Dnase treatment using the RNase-Free DNase Set (Qiagen, cat. no. 79254) and RNA was eluted in RNase-free water. cDNA was synthesised using M-MLV Reverse Transcriptase (Promega, cat. no. M1708). Quantitative PCR (qPCR) was performed using Power SYBR Green Master Mix (Thermo Fisher Scientific, cat. no. A25742) in 10 µl reactions, with forward and reverse primers each at a final concentration of 500 nM (Supplementary table 1). Expression of each target transcript was normalised to the *HPRT1* reference gene.

### pre-mRNA, mature mRNA and protein statistical comparisons

Technical replicate Ct values were averaged within each biological replicate. Each experiment was repeated in biologically independent experiments. Transcript abundance was calculated relative to housekeeping (ΔCt) and then converted to relative abundance (2^−ΔCt), such that increasing values represented increasing molecular abundance across all three datasets.

Western blot band intensities were quantified using ImageJ. Images were converted to 16-bit and inverted prior to analysis. Identically sized regions of interest were used to measure integrated density for each band, and background-corrected signal was calculated as integrated density − (background mean intensity × ROI area). Target protein abundance was normalised to the corresponding loading-control protein from the same sample, and normalised values were used as relative protein abundance for downstream statistical analyses.

For comparisons between two regulatory layers; pre-mRNA versus mature mRNA or mature mRNA versus protein, only cell lines with n=3 for both variables were included. Consequently, the identities and number of contributing cell lines differed between genes and comparisons. Associations between pre-mRNA and mature mRNA abundance, and between mature mRNA and relative protein abundance, were assessed using two-sided Pearson correlation. Statistical analyses and data visualisation were performed in R.

PLAYR probes and protocol were based on (36) and controls *HPRT1* Forward TTGCTTTCCTTGGTCAGGC, *HPRT1* Reverse TGGCTTCAGTTGTGACCCAG’

### ChIP-qPCR

HEK293T cells were grown in 10cm dishes to 95% confluence and crosslinked with 1% formaldehyde for 10min at room temperature. Cells were treated with 0.125M glycine for 5min, washed with ice cold PBS and centrifuged. Cell pellets were lysed in 50 mM Tris-HCl pH8.1, 1% SDS and 10mM EDTA with protease inhibitors, on ice for 10min. Chromatin was sheared by sonication, 20 cycles 15s on/30s off at 50% power and clarified by centrifugation. The chromatin was diluted tenfold in 1% triton x-100, 2mM EDTA, 150mM NaCl and 20 mM Tris-HCl pH 8.1.

Chromatin was pre-cleared with 20μL Protein G sepharose beads for 2h at 4°C with 0.5% BSA. The sample was rotated overnight at 4°C with 2μg of STRA8 antibody or IgG. Complexes were captured with 30μl of Protein G Sepharose for 1h and sequentially washed in low salt buffer, high salt buffer, LiCl buffer and twice with TE buffer. Elution of chromatin was performed in 1% SDS/0.1 M NaHCO₃ for 1 h at room temperature.

Crosslinks were reversed with 0.2M NaCl overnight at 65°C. Subsequently treated with 20μg proteinase K for 1h at 45°C and DNA was purified using the GeneJET PCR purification kit. Enrichment was quantified by qPCR using 3μl DNA (or equivalent 2% input). ChIP-qPCR primers were: *TEX12*, forward 5′-CACGAGTATTCCAAGTGG-3′ and reverse 5′-CAGCTCTCCCACCACCAC-3′; and *SYCE2*, forward 5′-GTGATCCGTCTGCCTCGGCC-3′ and reverse 5′-GGAGCACGCGTCACAACAC-3′.

### SC promoter activity, DNA methylation and chromatin accessibility

#### Promoter identification and cloning

Candidate proximal promoter regions of genes were selected based on the transcription start site and enrichment of factor binding sites. Selected regions were amplified from genomic DNA using Phusion High Fidelity DNA polymerase and cloned upstream of the luciferase reporter in vector pGL4.32 following double digestion with *Nhe*I/*BgI*I and ligation.

#### Luciferase reporter assays

HEK293T cells were seeded at 1.5 x 10^5 cells per well of a 6-well plate and transfected with promoter-luciferase pGL4.32 constructs with Lipofectamine 2000 following manufacturers directions. Where indicated, cells were treated with either DMSO or 1μM of all-trans retinoic acid (RA) 24h after transfection. 72h after transfection, cells were lysed and luciferase activity measured. B-galactosidase reporter plasmid was co-transfected as internal control.

#### *In vitro* promoter methylation

Promoter reporter plasmids were methylated in vitro using CpG methyltransferase *M.Sss*I. 1μg plasmid DNA and 160μM S-adenosyl-L-methionine were incubated with *M.Sss*I at 37°C for 1h, enzymes were then inactivated at 65°C for 20min. Constructs were transfected into HEK293T cells and promoter activity determined by luciferase assay. Where indicated, cells were treated with 1µM RA, 3µM 5-aza-2′-deoxycytidine (5-AZA), or both prior to harvesting.

#### RT-qPCR following epigenetic treatment

HEK293T cells were treated with 1µM RA, 3µM 5-AZA, 400nM trichostatin A (TSA), or combinations thereof. RNA was extracted using TRIzol and 1μg was reverse transcribed using M-MLV transcriptase and oligo (dT). qPCR was performed using SYBR green on a StepOne system with three technical repeats per sample. Primer sequences were: *TEX12*, forward 5′-CAGTGCCAGATAGTCCACAGCT-3′ and reverse 5′-GCTTTGCATAGGTAGACAACATTAG-3′; *SYCP3*, forward 5′-ACGAGAGCCTATGACTTTGAGAC-3′ and reverse 5′-CCCATATCTTCAACTACTCCTGC-3′; and *SYCE3*, forward 5′-GGACTTGGAAAAGCTATTAGAAGAG-3′ and reverse 5′-TGCAGTTGACGAAGGCATCCTC-3′. SC transcript abundance was normalised to *HPRT1* and relative expression was calculated using the (\(\mathcal(22)^{\mathcal{-\Delta \Delta Ct}}\) method.

#### Glucosylation qPCR methylation analysis

Endogenous promoter methylation was quantified in HEK293T, PC3 and U2OS cells using the EpiMark 5-hmC and 5-mC analysis kit. In short, genomic DNA was isolated with the DNeasy Blood and Tissue Kit according to manufacturers instructions and treated with T4 B-glucosyltransferase followed by digestion with *Msp*I or *Hpa*II. qPCR primers were promoter specific and contained at least one CCGG restriction site. *SYCP3*, forward 5′-GCGACGCTTCTGAGGCAAG-3′ and reverse 5′-GCGCGCAATCGCCCCATG-3′; *SYCE2*, forward 5′-CAAGCTCGCACCCTCTGCGC-3′ and reverse 5′-CAGTTGGCCGCGCAAGTTCTC-3′; and *SYCE3*, forward 5′-CTGCGGCGTCCGAGTCTG-3′ and reverse 5′-GACACAGCAGGCAAACGG-3′. A CCGG-free loading-control amplicon was amplified using forward 5′-TGAGCTCCAAGGAAACACAG-3′ and reverse 5′-GAAGAGCAGTCACTTGCCT-3′ primers. Glucosylated *Msp*I-resistant DNA was used to quantify 5hmC, while *Hpa*II resistance represented combined 5mC and 5hmC; 5mC abundance was calculated following subtraction of the 5hmC component. A CCGG-free amplicon was used as a loading control.

#### Statistical analysis

Data are presented as mean ± SD or SEM as indicated. Pairwise comparisons were performed using two-tailed unpaired Student’s t-tests where appropriate (normal distribution confirmed). Statistical significance was defined as p< 0.05.

#### Meiotic and post-meiotic protein abundance

Stage-resolved protein abundance was obtained from the mouse spermatogenesis proteomics dataset PXD017284 (37). Processed MaxLFQ values for *SYCP1, SYCP2, SYCP3, SYCE1, SYCE2, SYCE3, TEX12* and *SIX6OS1* were extracted for early leptotene/leptotene, zygotene, early, middle and late pachytene, and early and late diplotene. Replicate measurements were averaged within each stage. For the stage-resolved protein heatmap, values were log2-transformed and standardised independently for each protein using row-wise z-scores.

Post-meiotic protein dynamics were analysed using total-proteome data from PXD063123 (38). Protein abundance was extracted across spermatid Steps 1–2, 3–4, 5–6 and 13–14 and normalized within protein to the corresponding Step 1–2 value.

#### Meiotic RNA protein temporal analysis

Stage-resolved RNA abundance was obtained from the mouse spermatogenesis single-cell RNA-seq dataset GSE107644 (39) and compared with protein abundance from PXD017284 (37). Early and late diplotene protein measurements were combined to generate six matched meiotic stages: leptotene, zygotene, early, middle and late pachytene, and diplotene. RNA and protein profiles were expressed as log2 abundance relative to leptotene.

Gene-wise Pearson correlations were calculated between RNA and protein abundance at matched stages. Temporal offsets were assessed by shifting RNA abundance one, two or three stages earlier relative to protein, providing 6, 5, 4 and 3 matched stage pairs at lags 0–3, respectively. The same-stage analysis was additionally performed for housekeeping/control genes (*PGK1, PKM, LDHA, PSMC2, G6PDX, VCL, EIF4G1* and *HPRT*) and non-SC meiotic genes (*SPO11, MLH1, REC8, HORMAD1, HORMAD2, TEX11, TRIP13* and *TDRD1*).

#### MCF7 nascent transcription and promoter analysis

Cell-cycle-resolved nascent transcription was analysed using MCF7 GRO-seq dataset GSE94479 (40) in the hg19/GRCh37 coordinate system (41). Two replicates each for G0/G1, S and G2/M were averaged within cell cycle phase. Strand-specific GRO-seq signal was aligned to the selected transcription start site (TSS) and quantified from −2 kb to +5 kb relative to transcriptional orientation.

Promoter-proximal transcription was defined primarily as TSS to +150 bp and gene-body transcription as +500 bp to the transcription end site. Alternative promoter windows of −100 to +300 bp and −250 to +500 bp were also tested for robustness.

Reference transcript and TSS coordinates were obtained from Ensembl v75 (42). GRO-seq-supported candidate TSSs were compared with FANTOM5 CAGE data (43) and ENCODE/Riken MCF7 CAGE data (44). Candidate and canonical TSSs representing the same promoter region were collapsed, whereas spatially distinct promoter candidates were retained. *TEX12* was analysed using its Ensembl v75 reference promoter (42).

Bulk untreated-MCF7 H3K4me3, H3K27ac and RNA polymerase II ChIP-seq data were obtained from ENCODE (44). H3K4me3 was represented by two UW/Stamatoyannopoulos tracks (GSM945269), H3K27ac by the USC/Snyder dataset (GSM945854) and RNA polymerase II by the UT-A/Crawford dataset (GSM822295) (44). Signal was quantified in strand-oriented 50-bp bins across ±2 kb of each selected TSS using Megadepth (45). Tracks were normalized to their mean signal within ±1 kb of the active reference promoters *ACTB, GAPDH* and *RPLP0*, and replicate H3K4me3 profiles were averaged. CAGE and ChIP-seq datasets were used to support promoter identity and promoter state and were not interpreted as cell-cycle-resolved measurements.

#### Integrated analysis of meiotic SC gene regulation

Promoter DNA methylation was quantified from the GSE132446 COOL-seq dataset (46) using mm10/GRCm38 coordinates. Endogenous CpG methylation was calculated from WCG sites as methylated reads divided by total methylated and unmethylated reads, with counts summed across sites before calculation of the methylation fraction. Promoter methylation was quantified within ±1 kb of the selected TSS.

Chromatin accessibility was analysed using meiotic ATAC-seq GSE212117, histone modifications using GSE132446, nascent transcription using leChRO-seq GSE212118, and mature RNA abundance using poly(A)+ RNA-seq GSE212116 (26). Analyses were centred on leptotene/zygotene, pachytene and diplotene. H3K4me3 and H3K27ac were used as measures of promoter state and H3K36me3 as a measure associated with productive elongation. Promoter-proximal regulation was assessed using the ratio of promoter-proximal to gene-body leChRO signal; a reduction in this ratio accompanied by increased gene-body transcription was considered compatible with release into productive elongation.

Translation efficiency was determined in purified leptotene/zygotene spermatocytes using matched RNA-seq GSE307441 and Ribo-seq GSE307886 (47), calculated for each gene as Ribo-seq abundance divided by RNA-seq abundance. Translation-efficiency values were ranked only within the eight SC genes and represent a single L/Z measurement.

Protein timing was assessed using PXD017284 (37), together with RNA profiles from GSE107644 (39). Because these datasets were generated independently, RNA–protein comparisons were treated as descriptive temporal alignments.

#### ATAC-seq analysis for SC promoter accessibility (Figure 7n)

Publicly available ATAC-seq data from untreated HEK293T, PC3 and U2OS cells were obtained through ChIP-Atlas and visualised in IGV using the hg19 reference genome. Chromatin accessibility was examined across SC promoter regions. The datasets analysed were SRX14524940 (HEK293T), SRX10475867 (PC3) and SRX19611510 (U2OS), using the ChIP-Atlas significance threshold of 50 and q<1×10^-5.

##### Developmental and gonadal scRNA-seq

Processed human and mouse scRNA-seq datasets (GSE45719, GSE63818, GSE86146, GSE106487, GSE112013, GSE118127, GSE136441, GSE142585, GSE157329, GSE159883, GSE274603, E-MTAB-3929, E-MTAB-6967, E-MTAB-9388 and E-MTAB-10551) were harmonised by species and cell population. SC-gene symbols were standardised, including mapping *C14orf39* and 4930447C04Rik to *SIX6OS1*. Expression was reported as the percentage of cells with detectable transcripts and displayed on a 0–100% scale.

##### Multi-organ scRNA-seq

Human and mouse data were obtained from CZ CELLxGENE Census release 2025-11-08 and restricted to normal primary cells or nuclei. Raw counts were summarised by organ and harmonised cell type. Detection required ≥30 cells, ≥5% expressing cells and pooled expression ≥1 nTPM. Panels b–k show continuous log10(nTPM + 1) expression without the 5% cutoff.

---

**Supplementary Figure S1.**
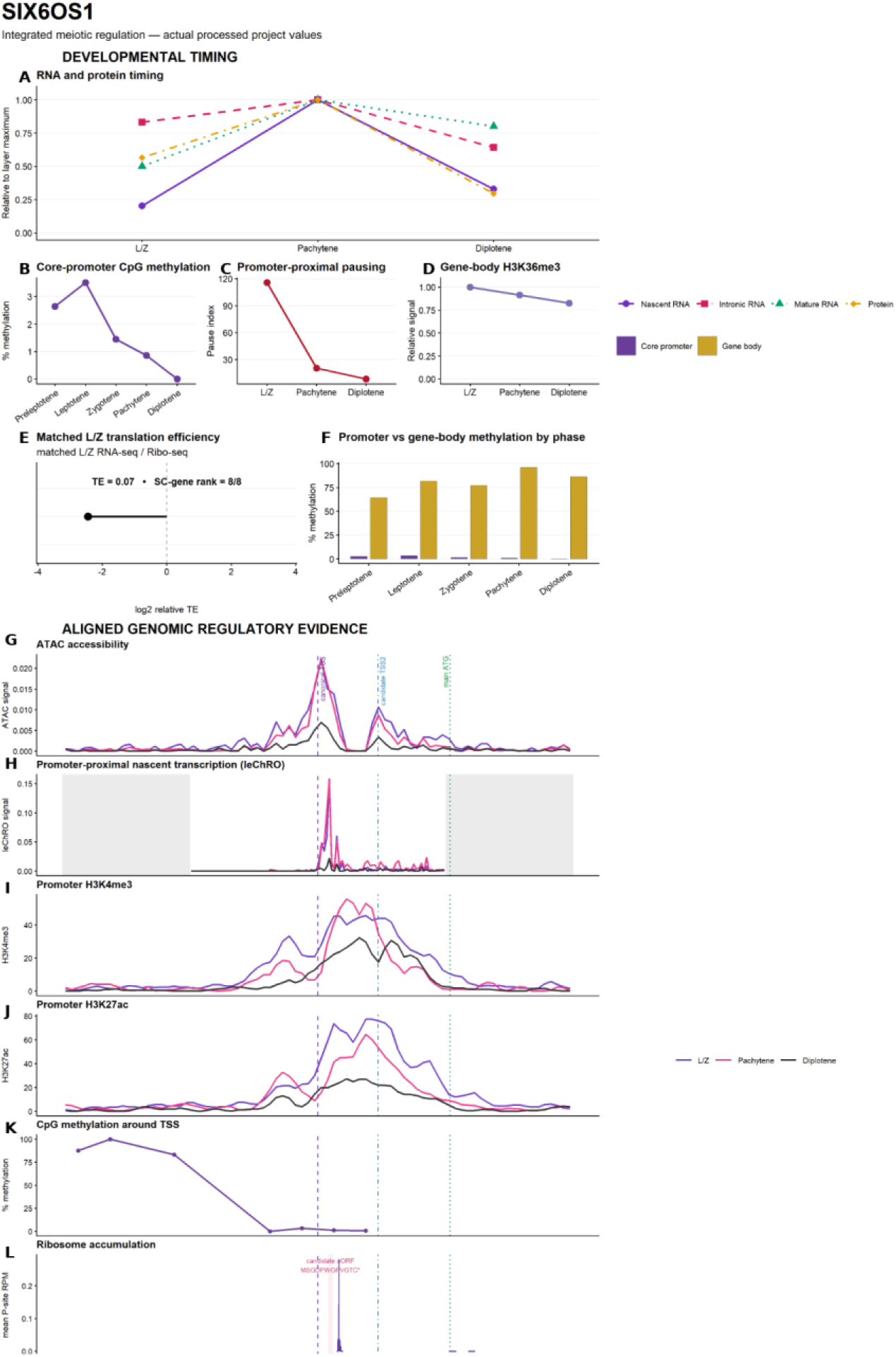
SIX6OS1 regulation combines a strong pause-release-compatible transition with low L/Z translation efficiency and a candidate 5’-leader gate. (A) Normalised nascent, intronic and mature RNA and protein abundance at L/Z, pachytene and diplotene. (B) Core-promoter CpG methylation. (C) Promoter-proximal pause index. (D) Relative gene-body H3K36me3. (E) Matched L/Z translation efficiency (TE) and rank among the eight SC genes. (F) Core-promoter versus gene-body CpG methylation. Aligned locus profiles show (G) ATAC-seq accessibility, (H) leChRO-seq nascent transcription, (I) H3K4me3, (J) H3K27ac, (K) CpG methylation and (L) ribosome-footprint density. Purple, pink and black profiles denote L/Z, pachytene and diplotene, respectively; dashed purple and dotted green lines mark the annotated TSS and main ATG. For SIX6OS1, additional vertical markers in panels G-L identify candidate TSS2 and the candidate uORF. *Data sources: GSE212118 (leChRO-seq), GSE21211c and GSE107c44 (RNA), GSE212117 (ATAC-seq), GSE13244c (histone marks), GSE30788c (positional ribosome footprints only), PXD017284 (protein), processed WGBS/RRBS values, and the final V18 matched L/Z RNA-seq/Ribo-seq TE dataset*.

**Supplementary Figure S2.**
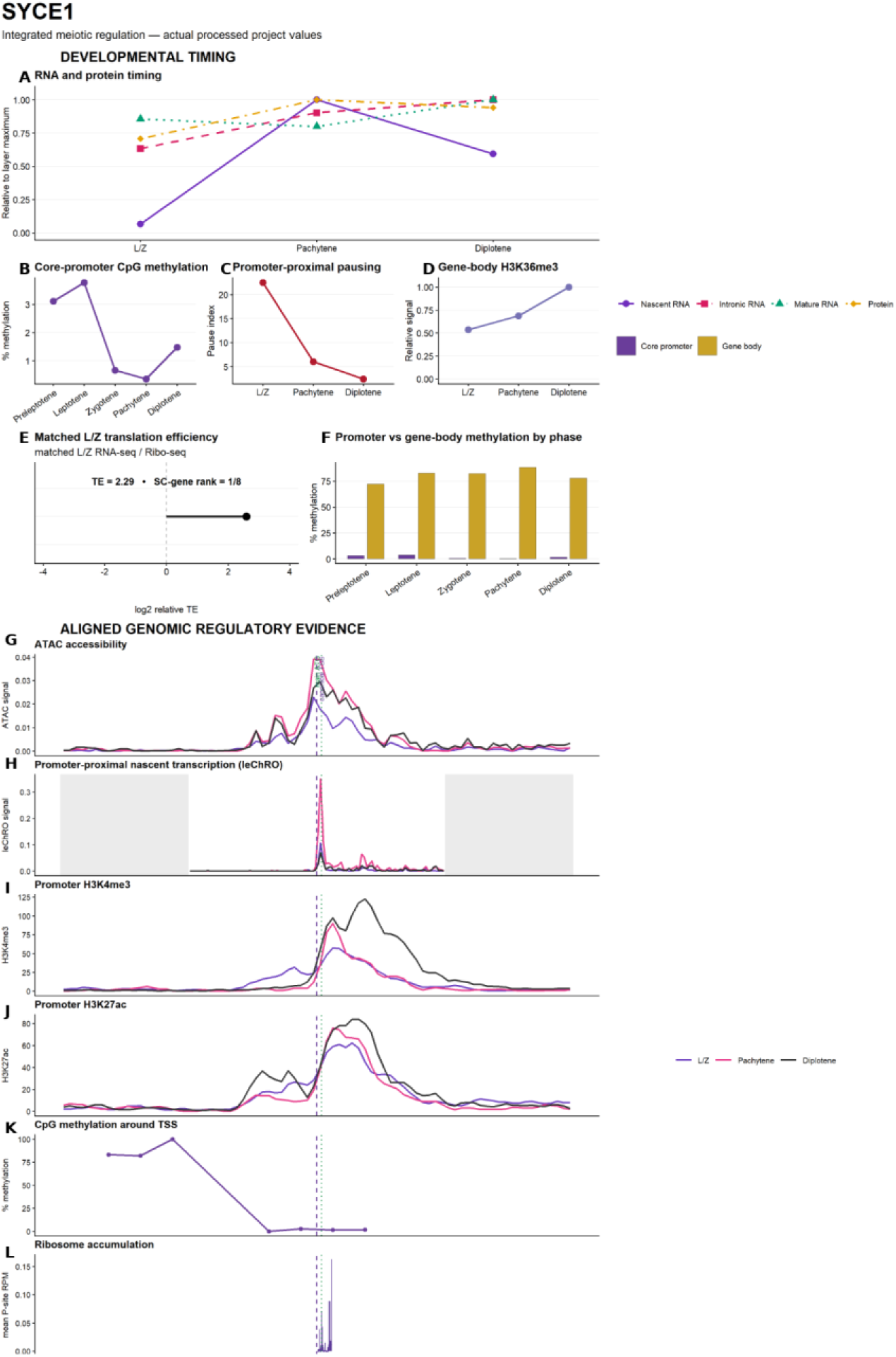
SYCE1 regulation is dominated by a pachytene pause-release-compatible transition followed by efficient L/Z translation. (A) Normalised nascent, intronic and mature RNA and protein abundance at L/Z, pachytene and diplotene. (B) Core-promoter CpG methylation. (C) Promoter-proximal pause index. (D) Relative gene-body H3K36me3. (E) Matched L/Z translation efficiency (TE) and rank among the eight SC genes. (F) Core-promoter versus gene-body CpG methylation. Aligned locus profiles show (G) ATAC-seq accessibility, (H) leChRO-seq nascent transcription, (I) H3K4me3, (J) H3K27ac, (K) CpG methylation and (L) ribosome-footprint density. Purple, pink and black profiles denote L/Z, pachytene and diplotene, respectively; dashed purple and dotted green lines mark the annotated TSS and main ATG. *Data sources: GSE212118 (leChRO-seq), GSE21211c and GSE107c44 (RNA), GSE212117 (ATAC-seq), GSE13244c (histone marks), GSE30788c (positional ribosome footprints only), PXD017284 (protein), processed WGBS/RRBS values, and the final V18 matched L/Z RNA-seq/Ribo-seq TE dataset*.

**Supplementary Figure S3.**
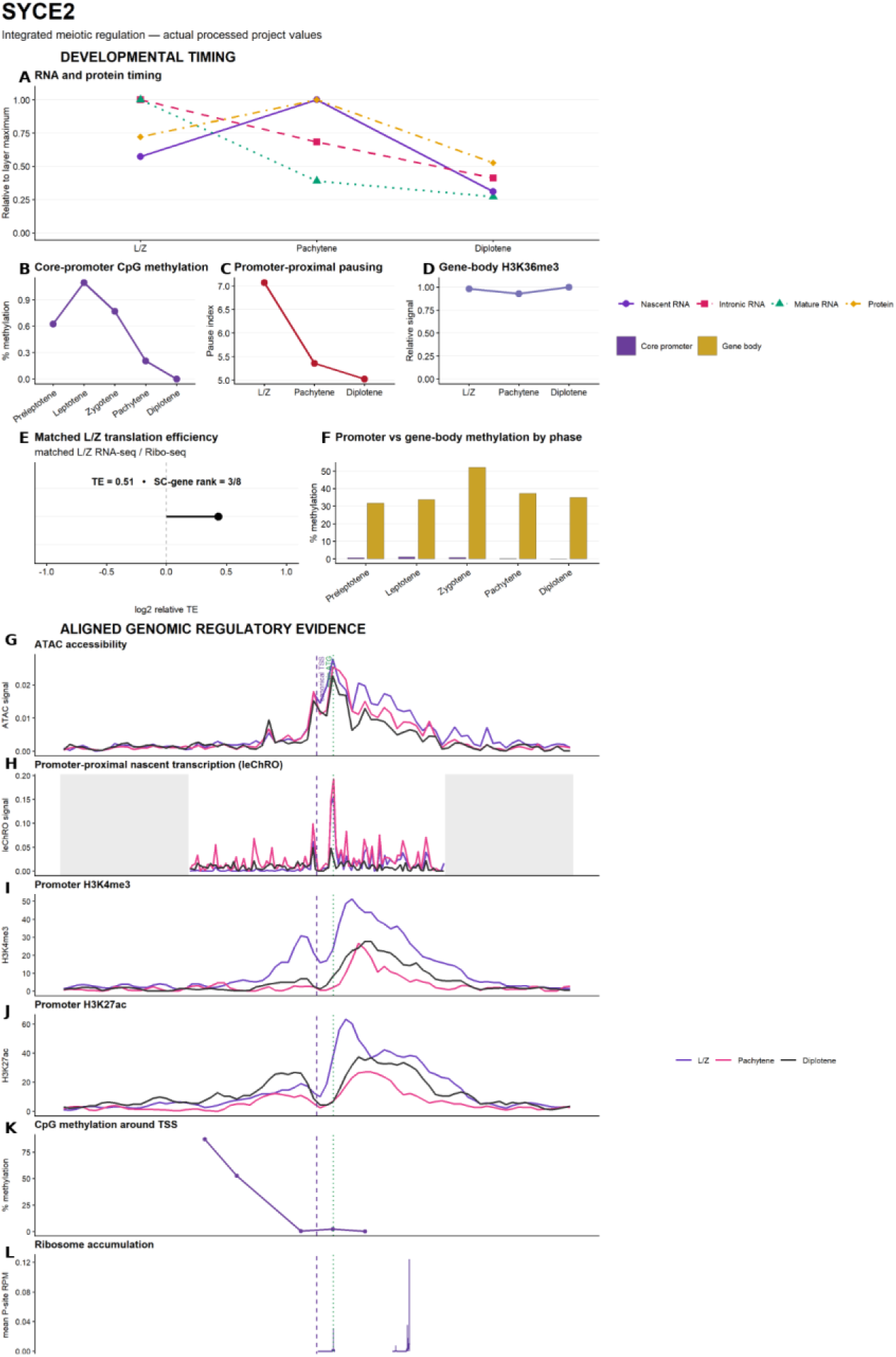
SYCE2 regulation is dominated by early promoter competence and RNA-pool persistence or turnover. (A) Normalised nascent, intronic and mature RNA and protein abundance at L/Z, pachytene and diplotene. (B) Core-promoter CpG methylation. (C) Promoter-proximal pause index. (D) Relative gene-body H3K36me3. (E) Matched L/Z translation efficiency (TE) and rank among the eight SC genes. (F) Core-promoter versus gene-body CpG methylation. Aligned locus profiles show (G) ATAC-seq accessibility, (H) leChRO-seq nascent transcription, (I) H3K4me3, (J) H3K27ac, (K) CpG methylation and (L) ribosome-footprint density. Purple, pink and black profiles denote L/Z, pachytene and diplotene, respectively; dashed purple and dotted green lines mark the annotated TSS and main ATG. *Data sources: GSE212118 (leChRO-seq), GSE21211c and GSE107c44 (RNA), GSE212117 (ATAC-seq), GSE13244c (histone marks), GSE30788c (positional ribosome footprints only), PXD017284 (protein), processed WGBS/RRBS values, and the final V18 matched L/Z RNA-seq/Ribo-seq TE dataset*.

**Supplementary Figure S4.**
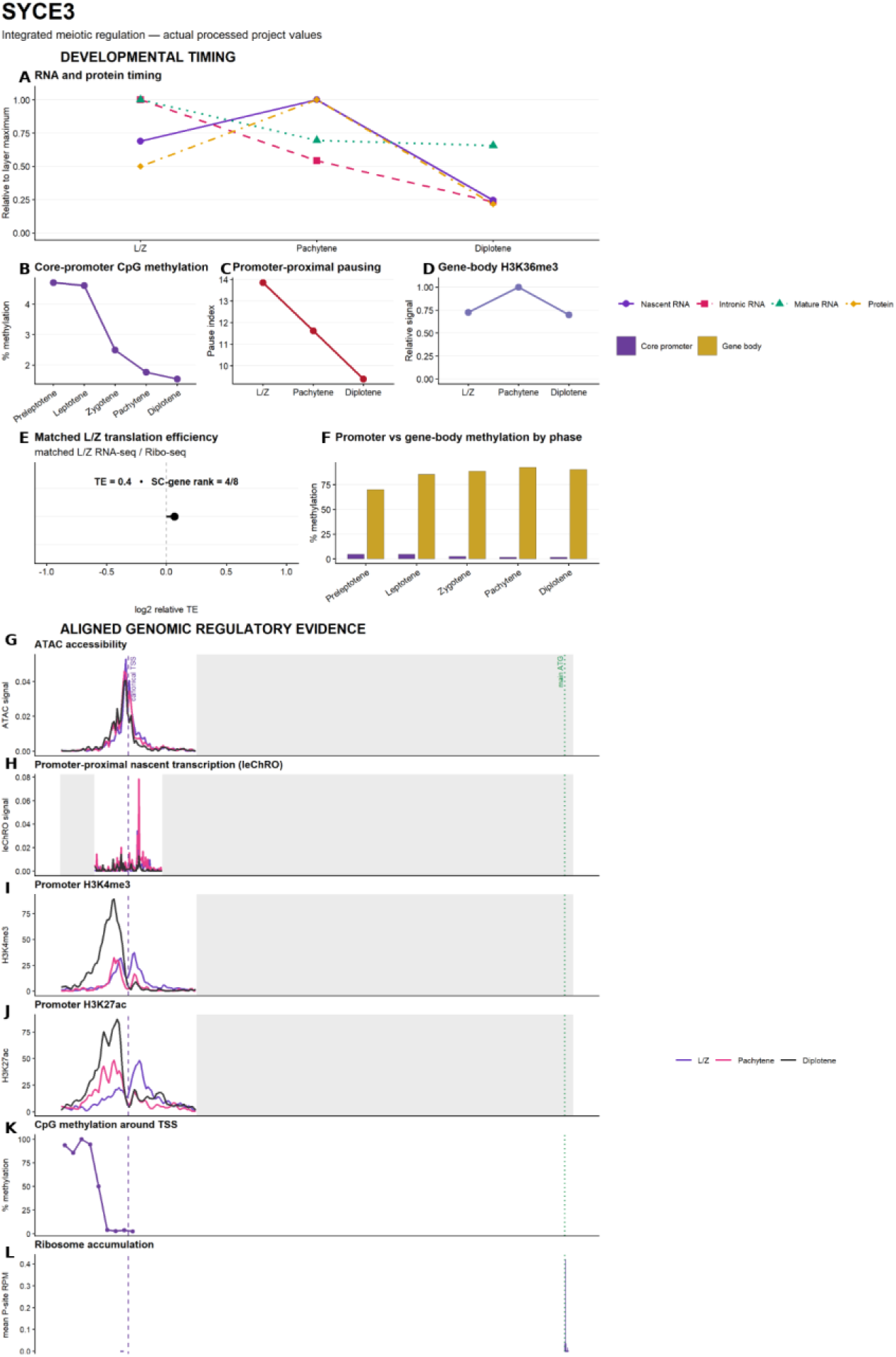
SYCE3 regulation is dominated by RNA persistence or turnover with a more modest promoter-proximal transition. (A) Normalised nascent, intronic and mature RNA and protein abundance at L/Z, pachytene and diplotene. (B) Core-promoter CpG methylation. (C) Promoter-proximal pause index. (D) Relative gene-body H3K36me3. (E) Matched L/Z translation efficiency (TE) and rank among the eight SC genes. (F) Core-promoter versus gene-body CpG methylation. Aligned locus profiles show (G) ATAC-seq accessibility, (H) leChRO-seq nascent transcription, (I) H3K4me3, (J) H3K27ac, (K) CpG methylation and (L) ribosome-footprint density. Purple, pink and black profiles denote L/Z, pachytene and diplotene, respectively; dashed purple and dotted green lines mark the annotated TSS and main ATG. *Data sources: GSE212118 (leChRO-seq), GSE21211c and GSE107c44 (RNA), GSE212117 (ATAC-seq), GSE13244c (histone marks), GSE30788c (positional ribosome footprints only), PXD017284 (protein), processed WGBS/RRBS values, and the final V18 matched L/Z RNA-seq/Ribo-seq TE dataset*.

**Supplementary Figure S5.**
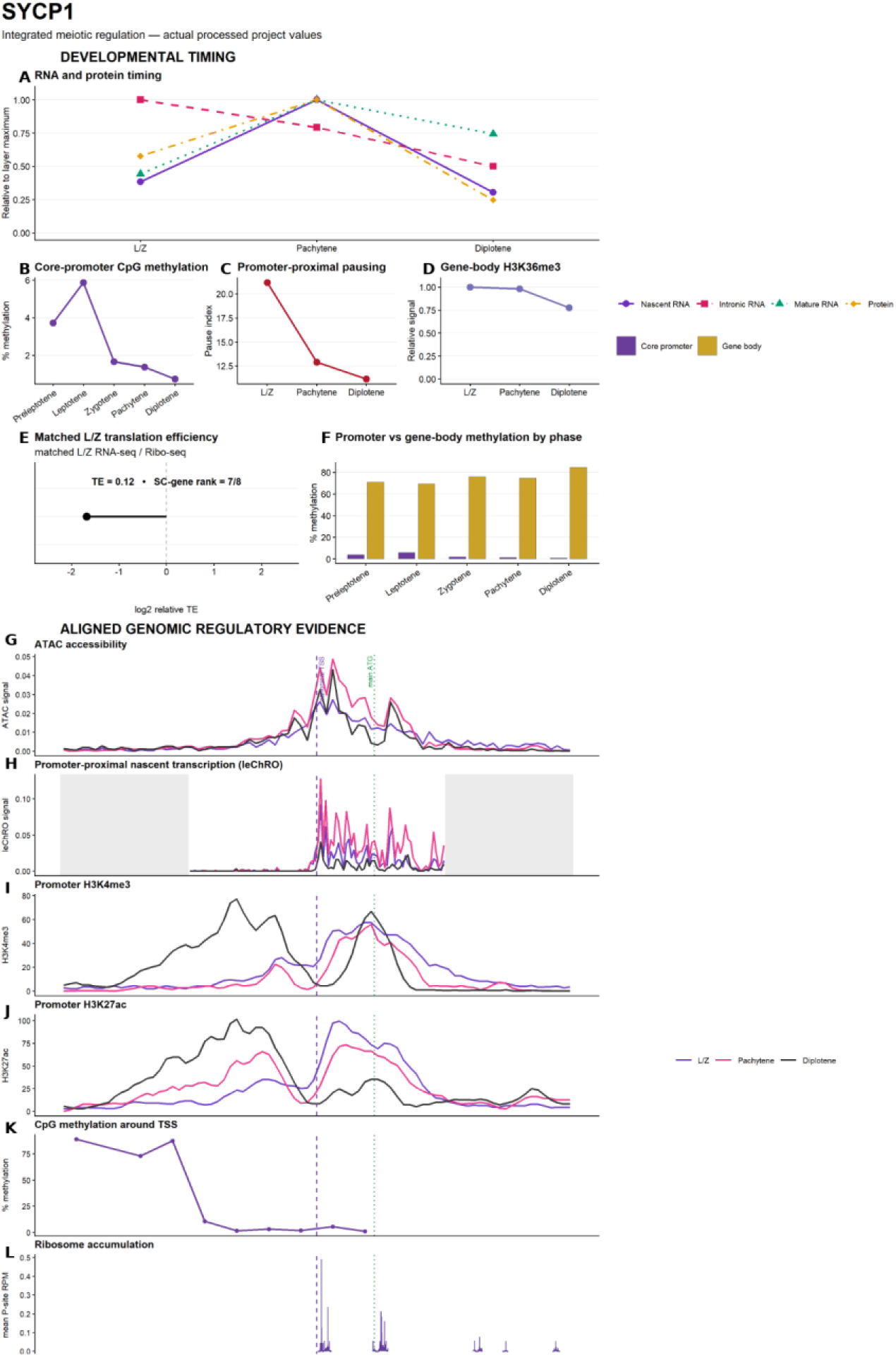
SYCP1 retains promoter-proximal regulation and shows a candidate downstream translation or protein-turnover gate. (A) Normalised nascent, intronic and mature RNA and protein abundance at L/Z, pachytene and diplotene. (B) Core-promoter CpG methylation. (C) Promoter-proximal pause index. (D) Relative gene-body H3K36me3. (E) Matched L/Z translation efficiency (TE) and rank among the eight SC genes. (F) Core-promoter versus gene-body CpG methylation. Aligned locus profiles show (G) ATAC-seq accessibility, (H) leChRO-seq nascent transcription, (I) H3K4me3, (J) H3K27ac, (K) CpG methylation and (L) ribosome-footprint density. Purple, pink and black profiles denote L/Z, pachytene and diplotene, respectively; dashed purple and dotted green lines mark the annotated TSS and main ATG. *Data sources: GSE212118 (leChRO-seq), GSE21211c and GSE107c44 (RNA), GSE212117 (ATAC-seq), GSE13244c (histone marks), GSE30788c (positional ribosome footprints only), PXD017284 (protein), processed WGBS/RRBS values, and the final V18 matched L/Z RNA-seq/Ribo-seq TE dataset*.

**Supplementary Figure S6.**
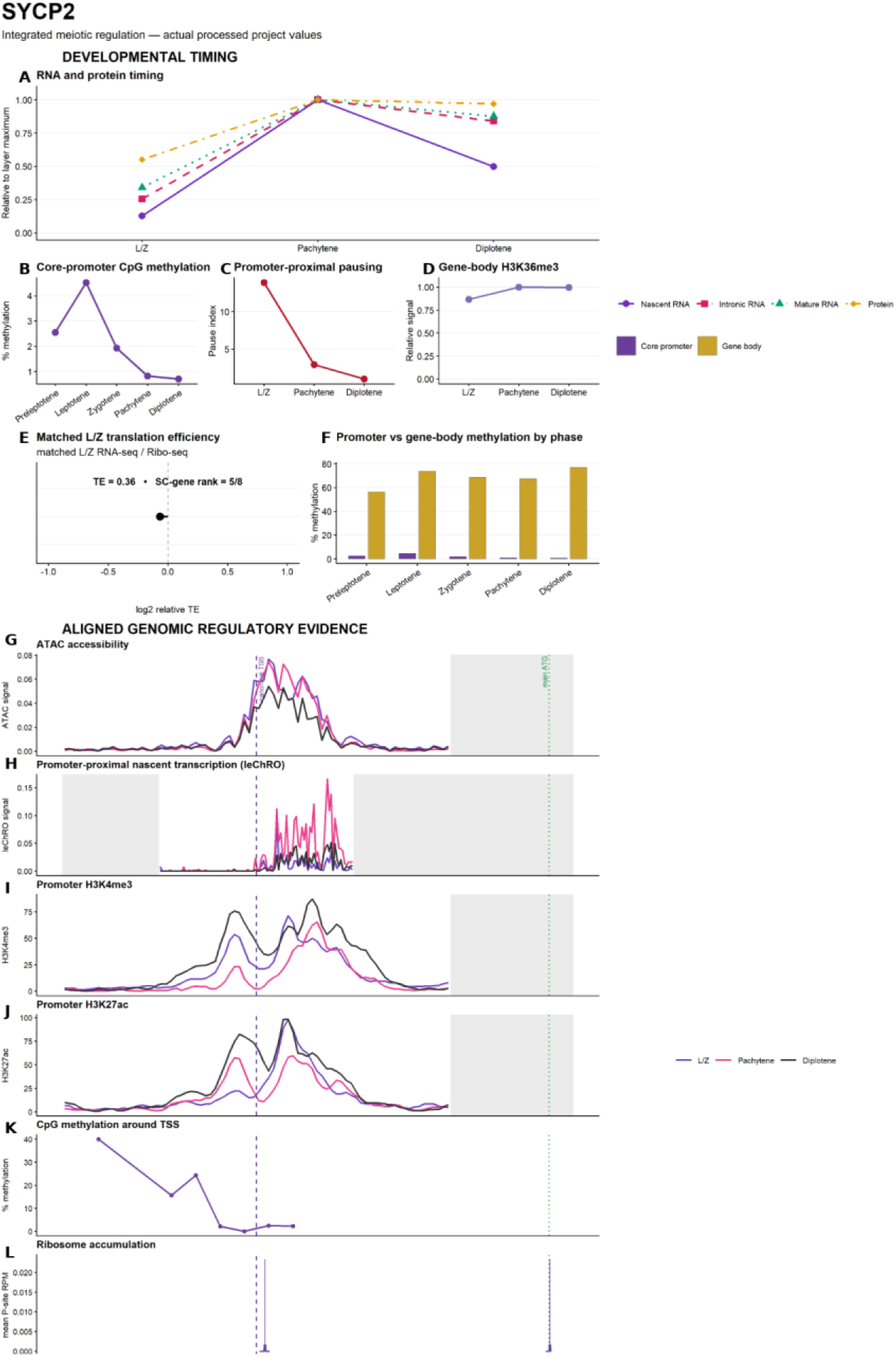
SYCP2 shows a strong pachytene pause-release-compatible transition into productive transcription. (A) Normalised nascent, intronic and mature RNA and protein abundance at L/Z, pachytene and diplotene. (B) Core-promoter CpG methylation. (C) Promoter-proximal pause index. (D) Relative gene-body H3K36me3. (E) Matched L/Z translation efficiency (TE) and rank among the eight SC genes. (F) Core-promoter versus gene-body CpG methylation. Aligned locus profiles show (G) ATAC-seq accessibility, (H) leChRO-seq nascent transcription, (I) H3K4me3, (J) H3K27ac, (K) CpG methylation and (L) ribosome-footprint density. Purple, pink and black profiles denote L/Z, pachytene and diplotene, respectively; dashed purple and dotted green lines mark the annotated TSS and main ATG. *Data sources: GSE212118 (leChRO-seq), GSE21211c and GSE107c44 (RNA), GSE212117 (ATAC-seq), GSE13244c (histone marks), GSE30788c (positional ribosome footprints only), PXD017284 (protein), processed WGBS/RRBS values, and the final V18 matched L/Z RNA-seq/Ribo-seq TE dataset*.

**Supplementary Figure S7.**
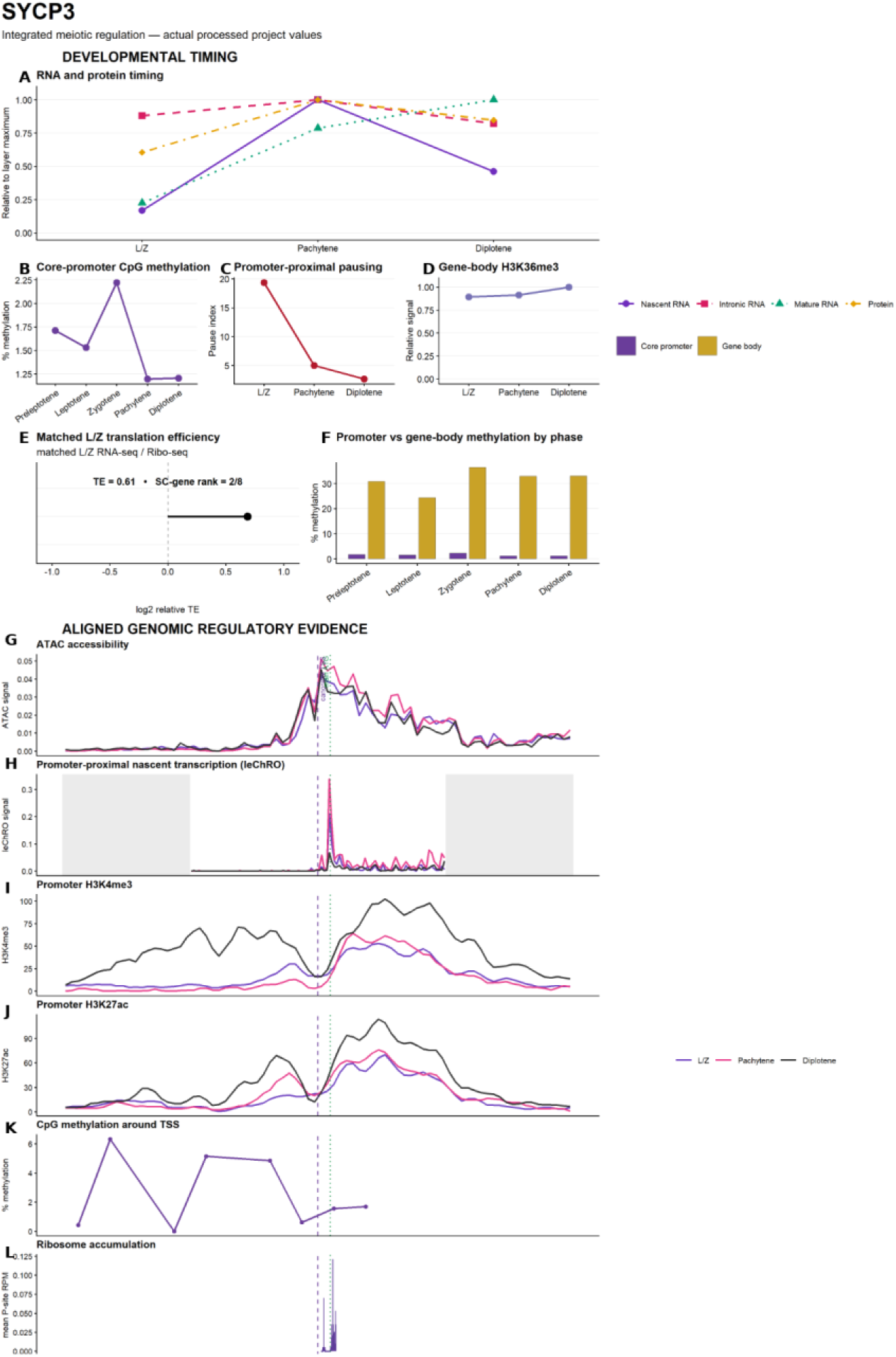
SYCP3 couples a pachytene pause-release-compatible transition to later mature-RNA accumulation and efficient L/Z translation. (A) Normalised nascent, intronic and mature RNA and protein abundance at L/Z, pachytene and diplotene. (B) Core-promoter CpG methylation. (C) Promoter-proximal pause index. (D) Relative gene-body H3K36me3. (E) Matched L/Z translation efficiency (TE) and rank among the eight SC genes. (F) Core-promoter versus gene-body CpG methylation. Aligned locus profiles show (G) ATAC-seq accessibility, (H) leChRO-seq nascent transcription, (I) H3K4me3, (J) H3K27ac, (K) CpG methylation and (L) ribosome-footprint density. Purple, pink and black profiles denote L/Z, pachytene and diplotene, respectively; dashed purple and dotted green lines mark the annotated TSS and main ATG. *Data sources: GSE212118 (leChRO-seq), GSE21211c and GSE107c44 (RNA), GSE212117 (ATAC-seq), GSE132446 (histone marks), GSE307886 (positional ribosome footprints only), PXD017284 (protein), processed WGBS/RRBS values, and the final V18 matched L/Z RNA-seq/Ribo-seq TE dataset*.

**Supplementary Figure S8.**
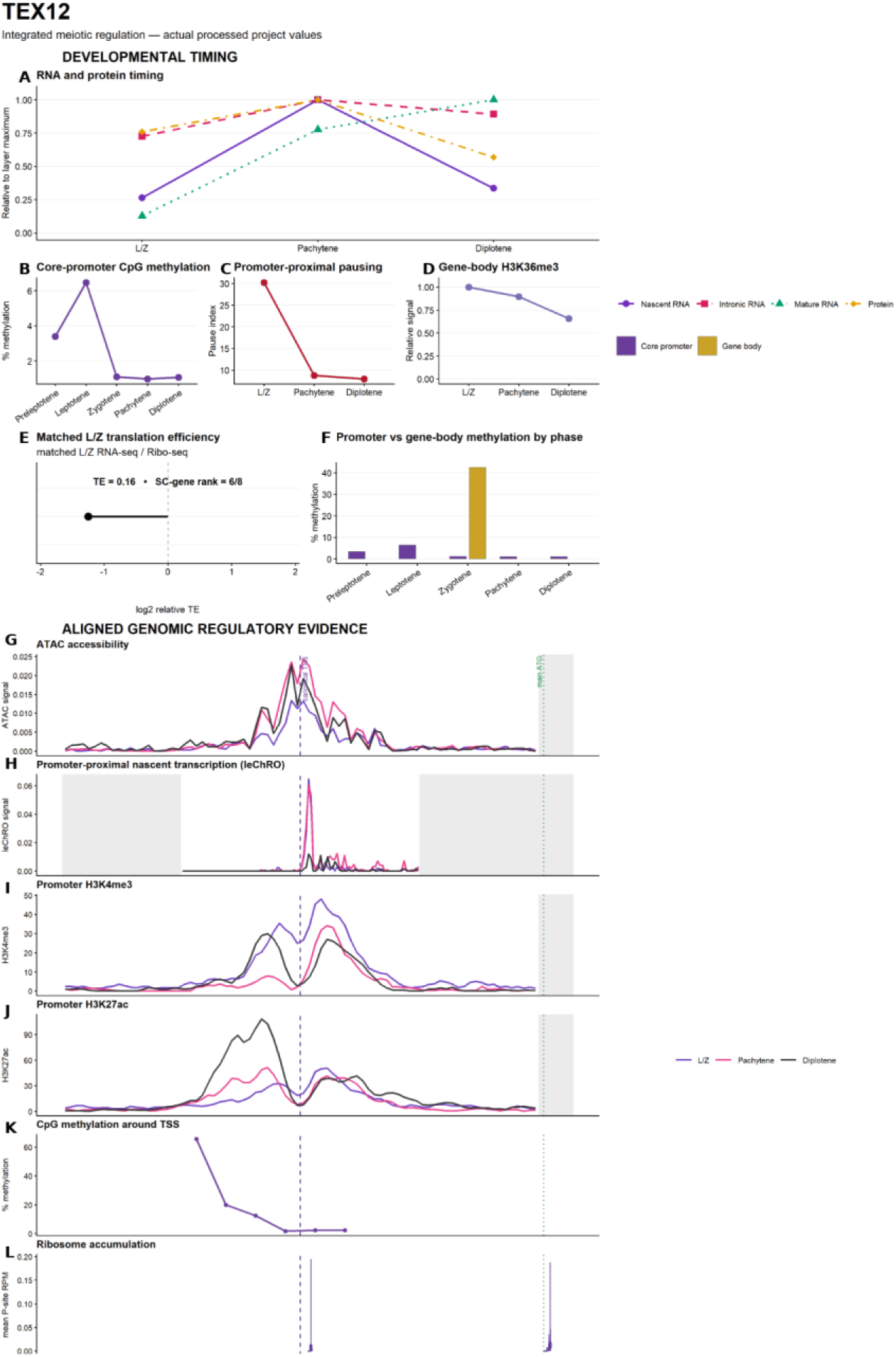
TEX12 couples a pachytene pause-release-compatible transition and later RNA accumulation to a candidate downstream gate. (A) Normalised nascent, intronic and mature RNA and protein abundance at L/Z, pachytene and diplotene. (B) Core-promoter CpG methylation. (C) Promoter-proximal pause index. (D) Relative gene-body H3K36me3. (E) Matched L/Z translation efficiency (TE) and rank among the eight SC genes. (F) Core-promoter versus gene-body CpG methylation. Aligned locus profiles show (G) ATAC-seq accessibility, (H) leChRO-seq nascent transcription, (I) H3K4me3, (J) H3K27ac, (K) CpG methylation and (L) ribosome-footprint density. Purple, pink and black profiles denote L/Z, pachytene and diplotene, respectively; dashed purple and dotted green lines mark the annotated TSS and main ATG. *Data sources: GSE212118 (leChRO-seq), GSE21211c and GSE107c44 (RNA), GSE212117 (ATAC-seq), GSE132446 (histone marks), GSE307886 (positional ribosome footprints only), PXD017284 (protein), processed WGBS/RRBS values, and the final V18 matched L/Z RNA-seq/Ribo-seq TE dataset*.

**Supplementary Figure S9.**
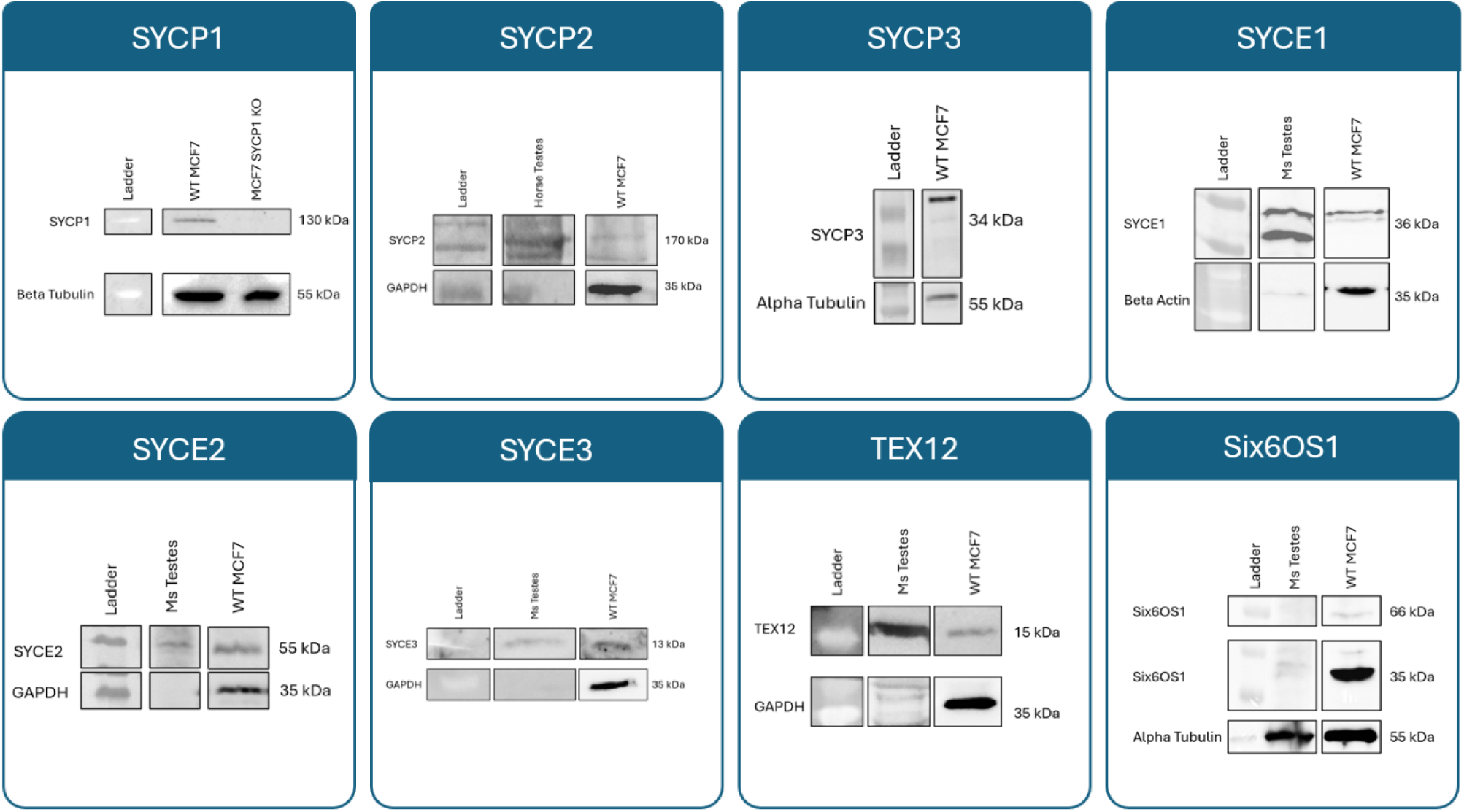
All eight synaptonemal complex proteins are expressed in breast cancer MCF7 cells. Whole-cell lysates from wild-type (WT) MCF7 cells were probed for each of the eight core SC proteins, alongside lysate of mouse (Ms) or horse testis and CRISPR KO control where available. GAPDH, Beta and Alpha Tubulin were used as loading control. Ladder lanes are included in each blot for molecular weight reference.

**Supplementary Table 1.**
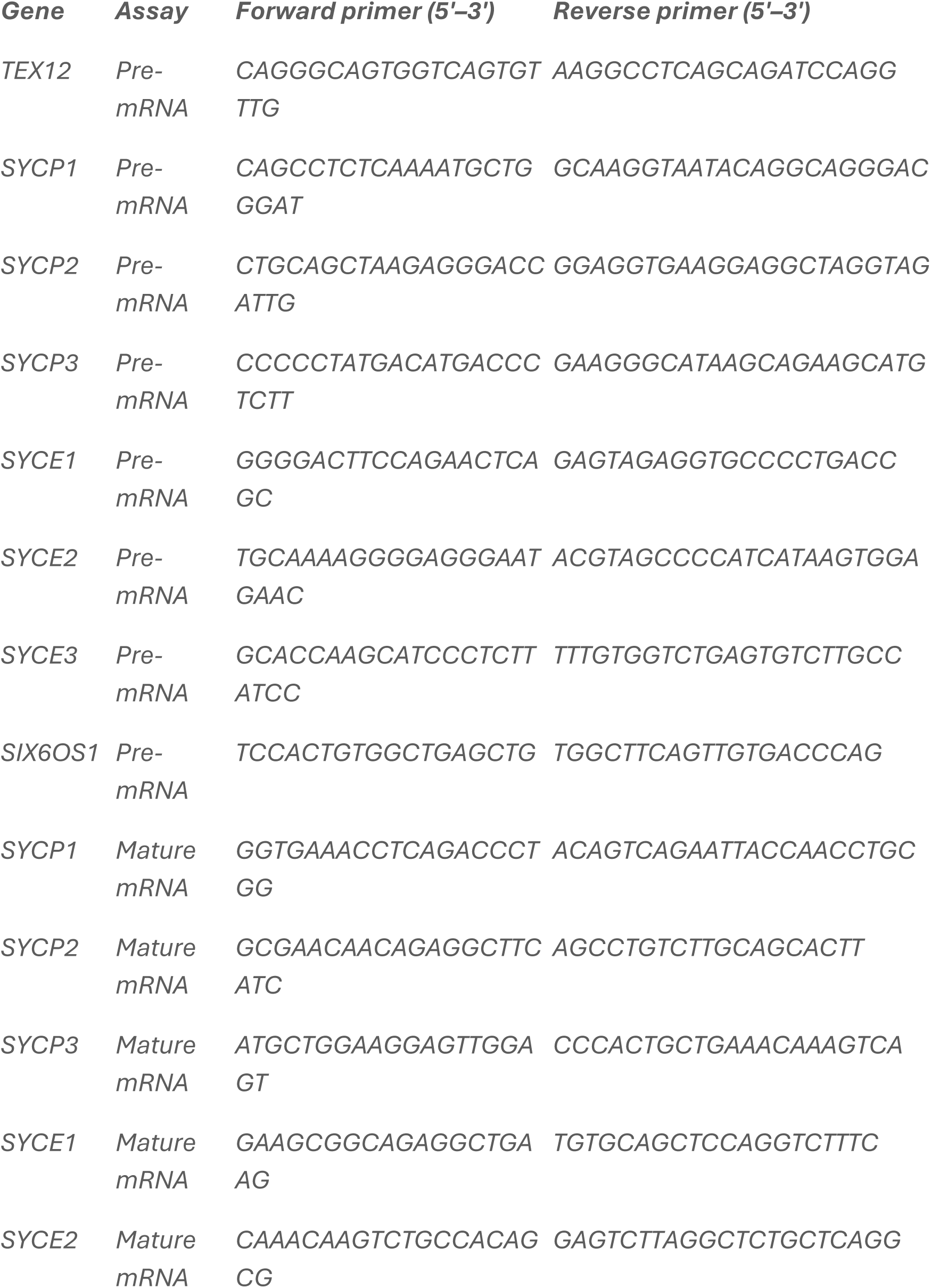
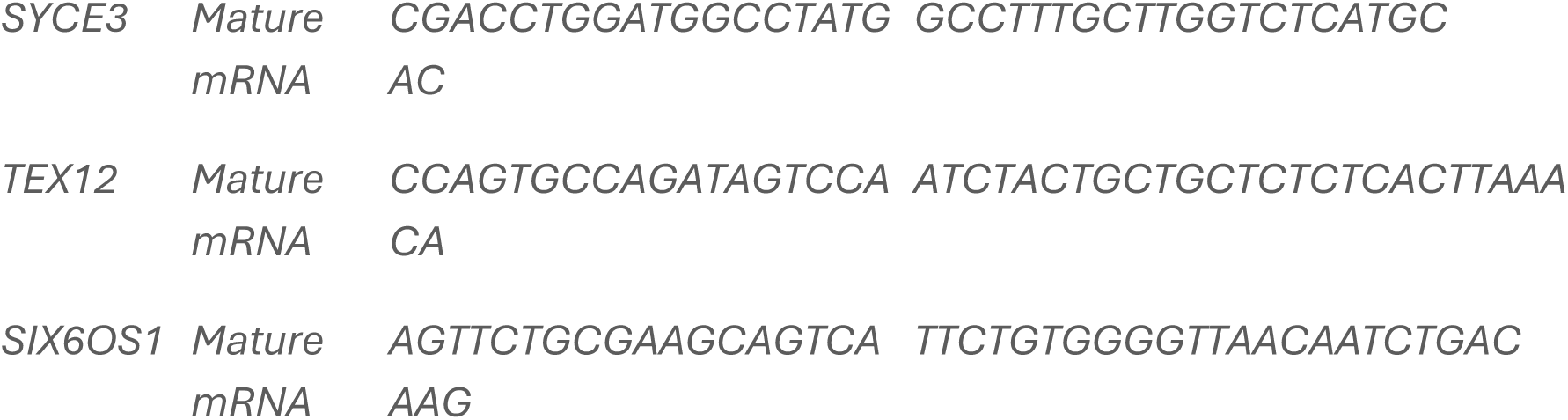
Human qPCR primers for detection of pre and mature mRNA.

**Supplementary Table 2.**
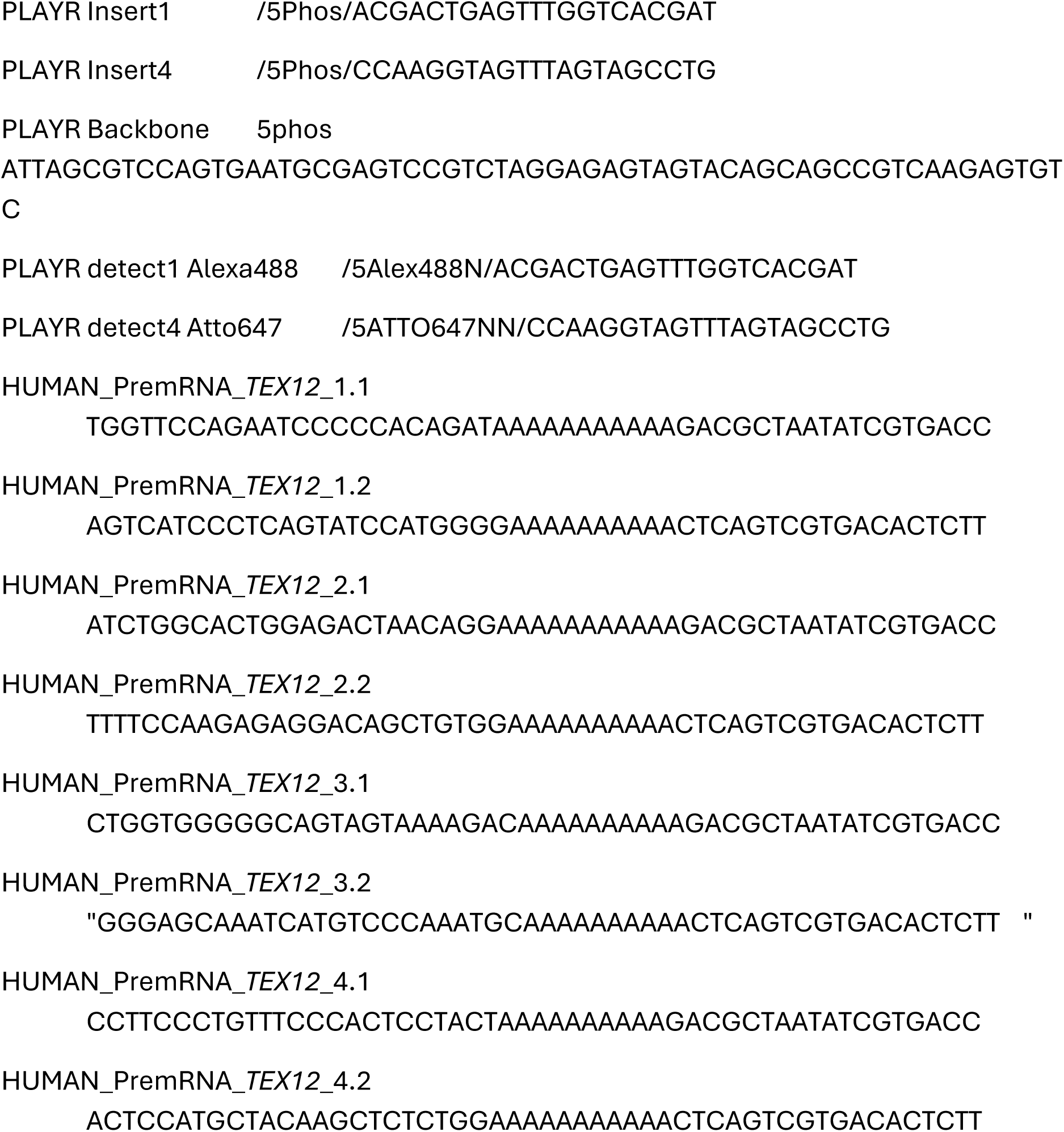
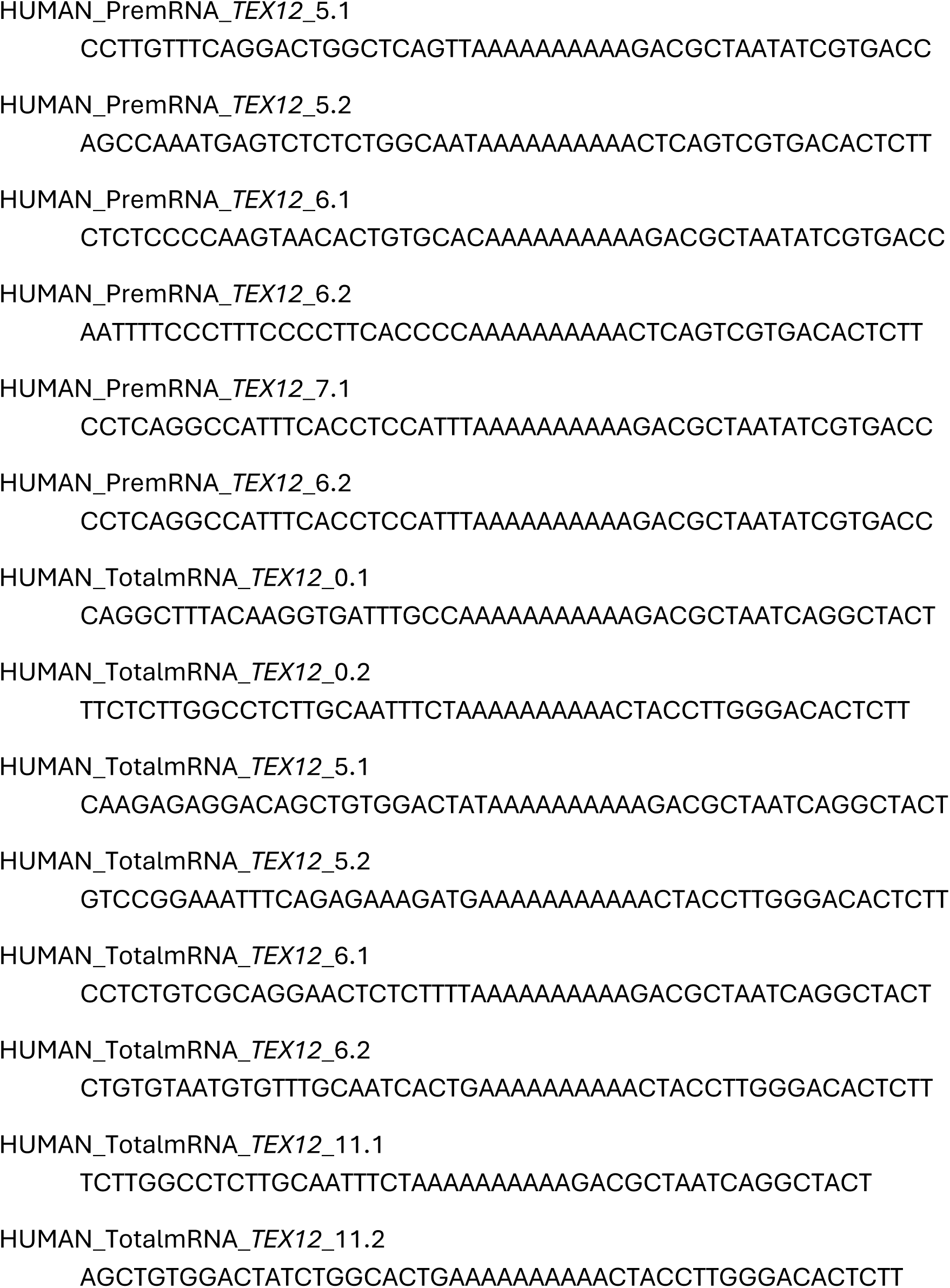
PLAYR probes for detection of pre and total *TEX12* mRNA.

## References

1. Fraune J, Schramm S, Alsheimer M, Benavente R. The mammalian synaptonemal complex: protein components, assembly and role in meiotic recombination. Exp Cell Res. 2012;318(12):1340–6.

2. Offenberg HH, Schalk JA, Meuwissen RL, van Aalderen M, Kester HA, Dietrich AJ, et al. SCP2: a major protein component of the axial elements of synaptonemal complexes of the rat. Nucleic acids research. 1998;26(11):2572–9.

3. Yuan L, Liu JG, Zhao J, Brundell E, Daneholt B, Höög C. The murine SCP3 gene is required for synaptonemal complex assembly, chromosome synapsis, and male fertility. Mol Cell. 2000;5(1):73–83.

4. Meuwissen RL, Offenberg HH, Dietrich AJ, Riesewijk A, van Iersel M, Heyting C. A coiled-coil related protein specific for synapsed regions of meiotic prophase chromosomes. The EMBO journal. 1992;11(13):5091–100.

5. Costa Y, Speed R, Ollinger R, Alsheimer M, Semple CA, Gautier P, et al. Two novel proteins recruited by synaptonemal complex protein 1 (SYCP1) are at the centre of meiosis. J Cell Sci. 2005;118(Pt 12):2755–62.

6. Schramm S, Fraune J, Naumann R, Hernandez-Hernandez A, Hoog C, Cooke HJ, et al. A novel mouse synaptonemal complex protein is essential for loading of central element proteins, recombination, and fertility. PLoS genetics. 2011;7(5):e1002088.

7. Hamer G, Gell K, Kouznetsova A, Novak I, Benavente R, Hoog C. Characterization of a novel meiosis-specific protein within the central element of the synaptonemal complex. Journal of cell science. 2006;119(Pt 19):4025–32.

8. Gomez HL, Felipe-Medina N, Sanchez-Martin M, Davies OR, Ramos I, Garcia-Tunon I, et al. C14ORF39/SIX6OS1 is a constituent of the synaptonemal complex and is essential for mouse fertility. Nat Commun. 2016;7:13298.

9. Argunhan B, Leung WK, Afshar N, Terentyev Y, Subramanian VV, Murayama Y, et al. Fundamental cell cycle kinases collaborate to ensure timely destruction of the synaptonemal complex during meiosis. The EMBO journal. 2017;36(17):2488–509.

10. Geisinger A, Benavente R. Mutations in Genes Coding for Synaptonemal Complex Proteins and Their Impact on Human Fertility. Cytogenetic and genome research. 2016;150(2):77–85.

11. De Smet C, Lurquin C, Lethe B, Martelange V, Boon T. DNA methylation is the primary silencing mechanism for a set of germ line- and tumor-specific genes with a CpG-rich promoter. Mol Cell Biol. 1999;19(11):7327–35.

12. Hackett JA, Sengupta R, Zylicz JJ, Murakami K, Lee C, Down TA, et al. Germline DNA demethylation dynamics and imprint erasure through 5-hydroxymethylcytosine. Science (New York, NY). 2013;339(6118):448–52.

13. Sou IF, Hamer G, Tee WW, Vader G, McClurg UL. Cancer and meiotic gene expression: Two sides of the same coin? Curr Top Dev Biol. 2023;151:43–68.

14. Brennan LC, Grinchuk OV, Pachon-Penalba M, Sou IF, Fawcett CJ, Nogueira CG, et al. Moonlighting role of meiotic SYCP1 in breast cancer: A chromatin-bound regulator of DNA repair, transcription, and drug resistance. Sci Adv. 2026;12(28):eaea2067.

15. Wang Y, Gao B, Zhang L, Wang X, Zhu X, Yang H, et al. Meiotic protein SYCP2 confers resistance to DNA-damaging agents through R-loop-mediated DNA repair. Nature communications. 2024;15(1):1568.

16. Hosoya N, Okajima M, Kinomura A, Fujii Y, Hiyama T, Sun J, et al. Synaptonemal complex protein SYCP3 impairs mitotic recombination by interfering with BRCA2. EMBO Rep. 2011;13(1):44–51.

17. Hosoya N, Ono M, Miyagawa K. Somatic role of SYCE2: an insulator that dissociates HP1α from H3K9me3 and potentiates DNA repair. Life Sci Alliance. 2018;1(3):e201800021.

18. Sandhu S, Sou IF, Hunter JE, Salmon L, Wilson CL, Perkins ND, et al. Centrosome dysfunction associated with somatic expression of the synaptonemal complex protein TEX12. Communications Biology. 2021;4(1):1371.

19. Tripathi N, Keshari S, Shahi P, Maurya P, Bhattacharjee A, Gupta K, et al. Human papillomavirus elevated genetic biomarker signature by statistical algorithm. J Cell Physiol. 2020;235(12):9922–32.

20. Duckworth AD, Gherardini PF, Sykorova M, Yasin F, Nolan GP, Slupsky JR, et al. Multiplexed profiling of RNA and protein expression signatures in individual cells using flow or mass cytometry. Nat Protoc. 2019;14(3):901–20.

21. Duckworth AD, Slupsky JR, Kalakonda N. Highly Multiplexed and Simultaneous Characterization of Protein and RNA in Single Cells by Flow or Mass Cytometry Platforms Using Proximity Ligation Assay for RNA. Methods in molecular biology (Clifton, NJ). 2024;2752:143–65.

22. Amouroux R, Nashun B, Shirane K, Nakagawa S, Hill PW, D’Souza Z, et al. De novo DNA methylation drives 5hmC accumulation in mouse zygotes. Nature cell biology. 2016;18(2):225–33.

23. Almatrafi A, Feichtinger J, Vernon EG, Escobar NG, Wakeman JA, Larcombe LD, et al. Identification of a class of human cancer germline genes with transcriptional silencing refractory to the hypomethylating drug 5-aza-2’-deoxycytidine. Oncoscience. 2014;1(11):745–50.

24. Almeida LG, Sakabe NJ, deOliveira AR, Silva MC, Mundstein AS, Cohen T, et al. CTdatabase: a knowledge-base of high-throughput and curated data on cancer-testis antigens. Nucleic Acids Res. 2009;37(Database issue):D816–9.

25. Coral S, Sigalotti L, Altomonte M, Engelsberg A, Colizzi F, Cattarossi I, et al. 5-aza-2’-deoxycytidine-induced expression of functional cancer testis antigens in human renal cell carcinoma: immunotherapeutic implications. Clin Cancer Res. 2002;8(8):2690–5.

26. Alexander AK, Rice EJ, Lujic J, Simon LE, Tanis S, Barshad G, et al. A-MYB and BRDT-dependent RNA Polymerase II pause release orchestrates transcriptional regulation in mammalian meiosis. Nature Communications. 2023;14(1):1753.

27. Kaye EG, Basavaraju K, Nelson GM, Zomer HD, Roy D, Joseph II, et al. RNA polymerase II pausing is essential during spermatogenesis for appropriate gene expression and completion of meiosis. Nature Communications. 2024;15(1):848.

28. Weber J, Salgaller M, Samid D, Johnson B, Herlyn M, Lassam N, et al. Expression of the MAGE-1 tumor antigen is up-regulated by the demethylating agent 5-aza-2’-deoxycytidine. Cancer Res. 1994;54(7):1766–71.

29. Zhang C, Kawakami T, Okada Y, Okamoto K. Distinctive epigenetic phenotype of cancer testis antigen genes among seminomatous and nonseminomatous testicular germ-cell tumors. Genes Chromosomes Cancer. 2005;43(1):104–12.

30. Litvinov IV, Cordeiro B, Huang Y, Zargham H, Pehr K, Doré MA, et al. Ectopic expression of cancer-testis antigens in cutaneous T-cell lymphoma patients. Clin Cancer Res. 2014;20(14):3799–808.

31. Niemeyer P, Türeci O, Eberle T, Graf N, Pfreundschuh M, Sahin U. Expression of serologically identified tumor antigens in acute leukemias. Leuk Res. 2003;27(7):655–60.

32. Shakib K, Norman JT, Fine LG, Brown LR, Godovac-Zimmermann J. Proteomics profiling of nuclear proteins for kidney fibroblasts suggests hypoxia, meiosis, and cancer may meet in the nucleus. Proteomics. 2005;5(11):2819–38.

33. Brennan LC, Grinchuk OV, Pachon-Penalba M, Sou IF, Fawcett CJ, Nogueira CG, et al. Moonlighting role of meiotic SYCP1 in breast cancer: A chromatin-bound regulator of DNA repair, transcription, and drug resistance. Science Advances. 2026;12(28):eaea2067.

34. Wang Y, Gao B, Zhang L, Wang X, Zhu X, Yang H, et al. Meiotic protein SYCP2 confers resistance to DNA-damaging agents through R-loop-mediated DNA repair. Nat Commun. 2024;15(1):1568.

35. Leiendecker L, Neumann T, Jung PS, Cronin SM, Steinacker TL, Schleiffer A, et al. Human Papillomavirus 42 Drives Digital Papillary Adenocarcinoma and Elicits a Germ Cell-like Program Conserved in HPV-Positive Cancers. Cancer Discov. 2023;13(1):70–84.

36. Duckworth AD, Gherardini PF, Sykorova M, Yasin F, Nolan GP, Slupsky JR, et al. Multiplexed profiling of RNA and protein expression signatures in individual cells using flow or mass cytometry. Nature Protocols. 2019;14(3):901–20.

37. Fang K, Li Q, Wei Y, Zhou C, Guo W, Shen J, et al. Prediction and Validation of Mouse Meiosis-Essential Genes Based on Spermatogenesis Proteome Dynamics. Mol Cell Proteomics. 2021;20:100014.

38. Zhu T, Zhu Y, Jiang X, Zhang X, Wang B, Chen Y, et al. Stage-Resolved Phosphoproteomic Landscape of Mouse Spermiogenesis Reveals Key Kinase Signaling in Sperm Morphogenesis. Adv Sci (Weinh). 2025;12(44):e08538.

39. Chen Y, Zheng Y, Gao Y, Lin Z, Yang S, Wang T, et al. Single-cell RNA-seq uncovers dynamic processes and critical regulators in mouse spermatogenesis. Cell Res. 2018;28(9):879–96.

40. Liu Y, Chen S, Wang S, Soares F, Fischer M, Meng F, et al. Transcriptional landscape of the human cell cycle. Proc Natl Acad Sci U S A. 2017;114(13):3473–8.

41. Church DM, Schneider VA, Graves T, Auger K, Cunningham F, Bouk N, et al. Modernizing reference genome assemblies. PLoS Biol. 2011;9(7):e1001091.

42. Flicek P, Amode MR, Barrell D, Beal K, Billis K, Brent S, et al. Ensembl 2014. Nucleic Acids Res. 2014;42(Database issue):D749–55.

43. Forrest AR, Kawaji H, Rehli M, Baillie JK, de Hoon MJ, Haberle V, et al. A promoter-level mammalian expression atlas. Nature. 2014;507(7493):462–70.

44. An integrated encyclopedia of DNA elements in the human genome. Nature. 2012;489(7414):57–74.

45. Wilks C, Ahmed O, Baker DN, Zhang D, Collado-Torres L, Langmead B. Megadepth: efficient coverage quantification for BigWigs and BAMs. Bioinformatics. 2021;37(18):3014–6.

46. Chen Y, Lyu R, Rong B, Zheng Y, Lin Z, Dai R, et al. Refined spatial temporal epigenomic profiling reveals intrinsic connection between PRDM9-mediated H3K4me3 and the fate of double-stranded breaks. Cell Res. 2020;30(3):256–68.

47. Xu J, Hu Y, Lu X, Cai Y, Li T, Gao M, et al. eEF1G Orchestrates Translation to Ensure Meiotic Progression in Transcriptionally Quiescent Spermatocytes. Adv Sci (Weinh). 2026:e76264.

